# Plant growth, biomass, and nitrogen utilization are impacted by mutations in C4 PEPCK1 in maize

**DOI:** 10.64898/2026.11.15.766785

**Authors:** Jennifer J. Arp, Maneesh Lingwan, Katherine M. Murphy, Jiahong Zhou, Patrick Z. Ellsworth, Anthony J. Studer, Shrikaar Kambhampati, Kevin L. Chu, Lauren M. Moore, Sarit Weissmann, Indrajit Kumar, Thomas P. Brutnell, Asaph B. Cousins, Todd C. Mockler, Stephen P. Moose, Doug K. Allen

**Affiliations:** Donald Danforth Plant Science Center, St. Louis, MO 63132 USA; United States Department of Agriculture, Agricultural Research Service, Sugarcane Unit, Houma, LA 70360 USA; Department of Crop Sciences, University of Illinois Urbana-Champaign, Urbana, IL 61801 USA; Virdis Genomics Consulting, Chesterfield, MO 63006 USA; School of Biological Sciences, Washington State University, Pullman, WA, 99164 USA; Bayer Crop Sciences, 800 N Lindbergh Blvd, Creve Coeur, Missouri 63141 USA; The Salk Institute for Biological Studies, 10010 North Torrey Pines Road, La Jolla, CA 92037 USA; KWS Gateway Research Center, 1005 N Warson Rd, St. Louis, MO 63132 USA

## Abstract

Efficient photosynthesis is crucial to agricultural productivity and can be constrained by the plant carbon-nitrogen balance, offering an opportunity for genetic improvement. Furthermore, soil N availability limits grain yields, and crops require substantial N fertilizer to be productive. In C_4_ plants, carbon-concentrating mechanisms respond to decreased soil nitrogen and share molecular intermediates with N metabolism. Maize, one of the most widely grown crops globally, uses a combination of two C_4_ photosynthetic pathways. Most carbon fixation occurs through the NADP-malic enzyme (NADP-ME) pathway, yet some fixation is also thought to occur via phospho*enol*pyruvate carboxykinase (PEPCK). However, genetic evidence for the contribution of PEPCK is lacking. Here, we used a transposon-based mutagenesis approach to study the function of the C4 isoform of PEPCK1 in maize. The C_4_ pathway intermediate aspartate accumulated to high concentrations in the leaves of the *pepck1-2::Ds* knock-down mutant, providing genetic support for the PEPCK secondary pathway as a complement to the dominant NADP-ME shuttle in maize. Field experiments over multiple seasons in either N-limiting or N-sufficient soils demonstrated the *pepck1*::*Ds* mutants produced higher vegetative and grain biomass without changes in leaf photosynthetic rates or parameters. This study confirms that PEPCK1 catalyzes the decarboxylation of carbon shuttled as aspartate, as an alternative route to decarboxylation by NADP-ME in a nominally NADP-ME subtype C_4_ species maize, providing clear genetic support for two-pathway operation. Further, the greater resilience and N-responsiveness of mutant plants are valuable traits for sustainable and subsistence farming in the future.

**One-sentence summary:** Transposon-derived maize *pepck1* mutant shows altered growth and productivity under altered soil nitrogen levels.

## Introduction

Approximately 500 million acres of maize are planted worldwide, yielding 1.1 billion tons of grain annually (Erenstein et al. 2022). Global average yields of 90-130 bushels per acre are strongly influenced by available nitrogen (N) and fertilizer applications; under optimal N conditions, productivity can exceed 220 bushels per acre (USDA-National Agricultural Statistics Service 2021). Despite its importance, N fertilizer is costly, energy-intensive to produce, and inaccessible to many farmers in low-income countries. Leaching and runoff of N contaminates groundwater, causes algal blooms and eutrophication, and contributes to N O emissions (Diaz and Rosenberg 2008). Improved germplasm with enhanced N use efficiency (NUE) would, therefore, have broad economic, environmental, food security, and ecological benefits (Menegat et al. 2022).

C_4_ photosynthesis has independently evolved at least 60 times in flowering plants (Sage 2004, 2016; Sage et al. 2011) and more than 20 times in grasses (Christin et al. 2013), concentrating CO around Rubisco’s active site. This suppresses photorespiration, improves carbon assimilation (Hatch and Slack 1966; Sage et al. 2011), and reduces the concentration of photosynthetic enzymes required per unit leaf area, resulting in greater NUE compared to C_3_ plants (Brown 1978). With few exceptions (Edwards et al. 2004), C_4_ photosynthesis requires the metabolic cooperation of two morphologically distinct cell types: mesophyll and bundle sheath (BS) cells. In the mesophyll, CO is converted to bicarbonate and combined with phosphoenolpyruvate (PEP) to generate oxaloacetate, which is reduced to malate or transaminated to aspartate (Figure 1). These transfer acids diffuse to the bundle sheath and are decarboxylated by NADP-malic enzyme (NADP-ME), NAD-malic enzyme (NAD-ME), or phosphoenolpyruvate carboxykinase (PEPCK), releasing CO and returning a 3-carbon molecule (pyruvate, PEP, or alanine) to the mesophyll to complete the C_4_ cycle (Bräutigam et al. 2014; Wang et al. 2014; Clapero et al. 2026a). Thus, C_4_ plants have been classified into three subtypes based on their primary bundle sheath decarboxylase: NADP-ME, NAD-ME, or PEPCK (Hatch et al. 1975). When aspartate serves as the transfer acid (NAD-ME and PEPCK pathways), an amino group is shuttled between cell types; N balance is maintained by generating alanine from pyruvate, as indicated by gene expression analysis and isotopic labeling (Gowik et al. 2011; Pick et al. 2011; Covshoff et al. 2016; Weissmann et al. 2016; Arrivault et al. 2017).

**Figure 1:**
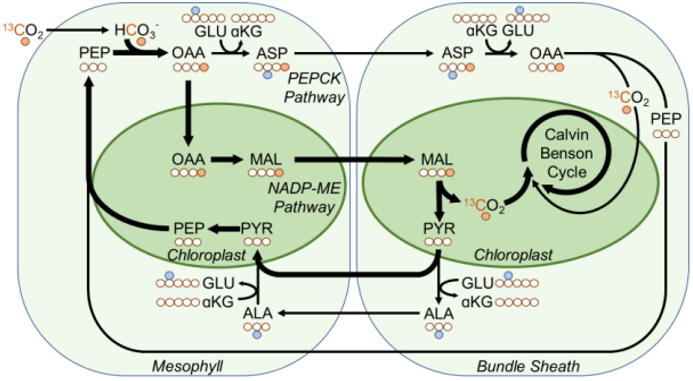
Topology of the C4 pathways in maize with the initial labeled carbon positions from ^13^CO_2_ incorporation highlighted in orange and nitrogen positions indicated in blue. Abbreviations: PEPCK: phosphoenolpyruvate carboxykinase, NADP-ME: NADP-malic enzyme, CO_2_: carbon dioxide, HCO_3_^-1^: bicarbonate, OAA: oxaloacetate, MAL: malate, ASP: aspartate, ASN: asparagine, PYR: pyruvate, PEP: phosphoenolpyruvate, ALA: alanine, CBC: Calvin-Benson Cycle.

Although functionally redundant in supplying CO□, the three pathways differ in their consequences for energy, reducing equivalents, N use, and metabolic gradients. Evidence suggests the pathways may operate in parallel and are more accurately described as two primary routes, NAD-ME and NADP-ME, with additional decarboxylation via PEPCK (Wang et al. 2014; Schlüter et al. 2019; Baccolini et al. 2026; Clapero et al. 2026b). Biochemical evidence suggests maize primarily uses the NADP-ME pathway, with PEPCK contributing an estimated 10–25% of decarboxylation activity (Weissmann et al. 2016; Arrivault et al. 2017). Early labeling experiments indicated that radiolabel from CO appeared in both malate and aspartate in maize (Chapman and Hatch 1979), and subsequent cellular analyses of PEPCK transcript, protein, enzyme activity, and isotopic labeling provide strong support for its role as an additional C_4_ decarboxylase (Furumoto et al. 1999; Wingler et al. 1999; Majeran et al. 2010; Pick et al. 2011; Weissmann et al. 2016; Arrivault et al. 2017), though direct genetic evidence has been lacking. Maintaining both pathways is thought to enable more flexible use of ATP and NADPH: the NADP-ME pathway shuttles CO alongside a reducing equivalent via malate, whereas PEPCK does not transfer reducing power, allowing the coordinated use of both pathways to balance CO and NADPH supply in BS cells in response to metabolic demand (Furbank 2011; Stitt and Zhu 2014; Wang et al. 2014; Clapero et al. 2026a).

Nitrogen is central to regulating both C_3_ and C_4_ photosynthesis and is allocated differently across species. In maize leaves, photosystem I proteins account for twice the proportion of leaf N as in rice (Makino et al. 2003), while Rubisco constitutes only 5–9% of total leaf N (Sage et al. 1987). This contrasts markedly with C_3_ plants, where Rubisco alone accounts for more than half of soluble protein and roughly 25% of total leaf N (Spreitzer and Salvucci 2002; Makino et al. 2003). Rubisco evolved in C_4_ plants for faster kinetics and higher CO assimilation rates enabled by the CO -enriched environment created by the carbon-concentrating mechanism (Yeoh et al. 1980; Seemann et al. 1984; Sage 2002), and the resulting reduction in Rubisco investment is a primary driver of enhanced NUE in C_4_ plants (Ghannoum et al. 2011). Among C_4_ subtypes, NUE also varies: NADP-ME and PEPCK species exhibit higher NUE than NAD-ME grasses (Ghannoum et al. 2005; Pinto et al. 2014, 2016), reflecting differences in N partitioning to the thylakoid or other tissues. C_4_ pathway enzyme levels are highly sensitive to plant N status (Sugiharto et al. 1990) and influence photosynthetic rate under non-limiting light (Matsuoka et al. 2001), further reflecting the interconnectivity of C_4_ and nitrogen pathways. Unlike NADP-ME, the PEPCK pathway shuttles amino acids and shares enzymes with N metabolism, directly linking carbon and N metabolism in the leaf (Majeran et al. 2005). Aspartate, in particular, connects N assimilation, protein synthesis, and the TCA cycle with C_4_ photosynthesis (Azevedo et al. 2006; Galili 2011) and contributes to improved NUE in certain NADP-ME species (Majeran et al. 2005; Bräutigam and Gowik 2016). Under N limitation, maize reduces aspartate shuttling while maintaining malate flux, shifting toward greater reliance on NADP-ME (Khamis et al. 1992).

Despite this biochemical understanding, no genetic mutants exist to test the overlapping roles of the C_4_ pathways in maize. We hypothesized that the NADP-ME and PEPCK pathways differ in how they coordinate with N metabolism. To assess the contribution of PEPCK in photosynthesis and N metabolism, we used transposon-based mutagenesis, employing a two-step transposition screen with a donor Dissociator (Ds) element located 600 kb away to reduce the decarboxylation activity of the primary C4 isoform PEPCK1. Carbon partitioning by isotopic labeling, metabolic pool sizes, plant growth, yield, and photosynthetic traits were assessed in homozygous mutants and near-isogenic W22 inbred parents under controlled greenhouse conditions and through multiple seasons of field trials. Our findings provide genetic, biochemical, and physiological evidence in support of an active PEPCK pathway in maize based most directly on changes in aspartate and alanine pool sizes, and increased biomass. These studies offer new insight into optimizing yields under both N-limited and N-replete conditions for the benefit of sustainable and subsistence farming.

## Results

### PEPCK1 mutants were generated using Ac/Ds

Transposon mutagenesis targeted the phosphoenolpyruvate carboxykinase1 (PEPCK1) gene (Zm00004b001002) to examine the contribution of the PEPCK pathway to maize performance. *Ds* preferentially inserts into closely linked sequences (McCLINTOCK 1950), so a two-step procedure was used to first mobilize the closest existing *Ds* donor, approximately 600kb from PEPCK1, to a site physically closer to the PEPCK1 target locus in the W22 inbred (Ahern et al. 2009; Kolbe et al. 2018). In the first screen, four genes in the intervening sequence between PEPCK1 and the donor *Ds* were assayed for insertions, along with PEPCK1. No insertions were recovered in PEPCK1; however, a single insertion allele was recovered in one of the intervening genes (Zm00004b001005), located 140kb away from the PEPCK1 locus. This new insertion allele was then used to generate a second population from which two insertion alleles were identified in PEPCK1. The precise location of the insertion alleles was determined by sequencing. The first insertion (*pepck1-m1::Ds*, hereafter referred to as *pepck1-1*) is located at the intron7-exon8 junction, and the second insertion (*pepck1-m2*::*Ds*, hereafter referred to as *pepck1-2*) is within the 3’UTR (Figure 2A). Plants hemizygous for the insertion allele were self-pollinated to generate true-breeding F_2_ populations that no longer carried the source of *Ac* transposase. Because all material was maintained in the W22 inbred and utilized an Ac/Ds transposon approach, near-isogenic comparisons could be made between the homozygous insertion alleles and wild-type parental lines. Assaying mRNA expression by qPCR of mature leaf tissue from the V8 growth stage showed *pepck1-1* expression was reduced to 61% of wild-type levels, whereas *pepck1-2* was further decreased to 36% (Figure 2B).

**Figure 2:**
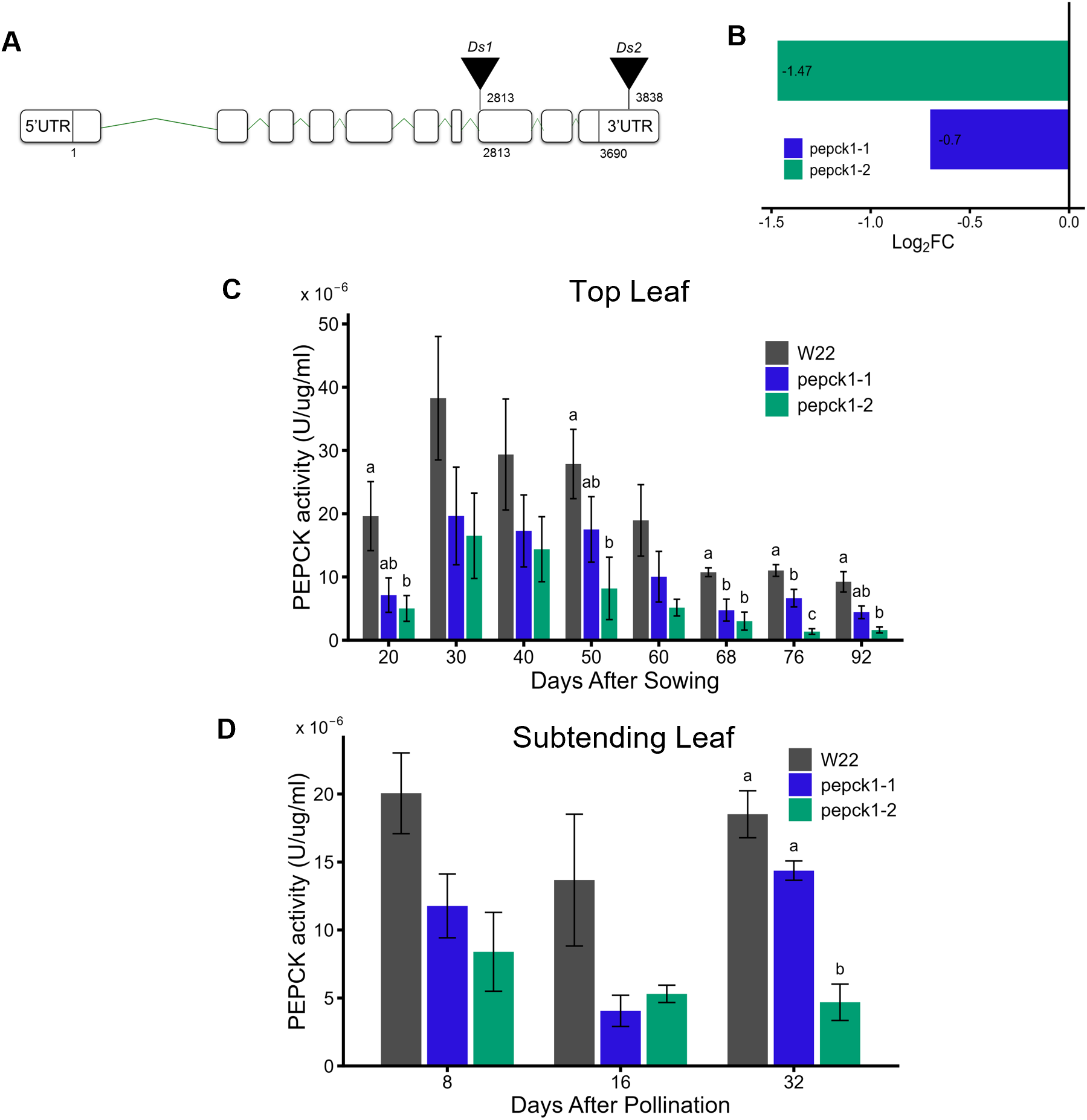
A.) Location of the *Ds* transposon in the two *pepck::Ds* alleles. *pepck1-1* is located at the junction of intron 7 and exon 8 at base 2813; *pepck1-2* is in the 3’UTR at base 3838. B.) Reduced gene expression of *pepck1-1* and *pepck1-2* determined by qPCR. PEPCK enzyme activity in *pepck1-1* and *pepck1-2* in top C.) and subtending D.) leaves. Error bars represent means ±standard error. Compact letter assignment represents significant differences (p < 0.05) among the three genotypes. No letter is displayed if there is no difference. (n = 6 in C and n = 3 in D, ANOVA followed by Tukey’s multiple comparisons test).

### PEPCK enzyme activity was reduced in mutants assayed over development

The effects of the mutations were assessed by measuring PEPCK activity throughout the plant life cycle. The top leaf was harvested at ten-day intervals during plant development for each line (Figure 2C) and after pollination the top and subtending leaves were harvested at 8, 16, and 32 days (Figure 2D); leaves were assayed according to (Sharwood et al. 2014, 2016) as described in the methods. As an assay control, PEPCK activity was monitored by varying the amount of PEP substrate; activity responded linearly, as expected, to changes in substrate or enzyme levels (Suppl. Figure 1A). Measured activities for W22 were comparable to prior reports for other species (Sharwood et al. 2016). PEPCK enzyme activity in the mutants was significantly lower than in W22 at all developmental points tested, with a larger decrease in *pepck1-2*, consistent with expression differences (Figure 2B-D). In independent top and subtending leaf comparisons, PEPCK activity was always significantly lower in both mutants compared to W22 (Suppl. Figure 1B)

### Aspartate accumulated as a result of the *pepck1* mutation

In *pepck1-2*, aspartate concentrations were higher than W22 (Figure 3A). The aspartate pool size was 3.5-fold greater in *pepck1-2* than in W22 at high N and 3.7-fold greater at low N, suggesting that reduced PEPCK activity led to aspartate accumulation. The aspartate pool size was also highly responsive to N. In *pepck1-2* at high N, aspartate accumulated to 6668 nmol g^-1^ FW, compared to 2680 nmol g^-1^ FW at low N. The malate pool was larger than the aspartate pool in W22, although not in the mutant (Figure 3A). Alanine and pyruvate in *pepck1-2* plants were reduced, consistent with decreased PEPCK activity and nitrogen shuttling through a combined PEPCK/NADP-ME pathway. Alanine pools were 1307±117 (standard error) nmol g^-1^ FW in the mutant at high N and 896±107 nmol g^-1^ FW at low N, whereas in W22 the pools were larger, being 2166±493 nmol g^-1^ FW and 1004±52 nmol g^-1^ FW at high and low N, respectively (Suppl. Figure 2). Pool sizes for metabolites involved in C_3_ photosynthesis pathways were slightly smaller in *pepck1-2* (Suppl. Figure 2). Changes in aspartate and alanine in genetic mutants support a role for the PEPCK pathway in maize and warranted further testing to gauge it relative to NADP-ME.

**Figure 3:**
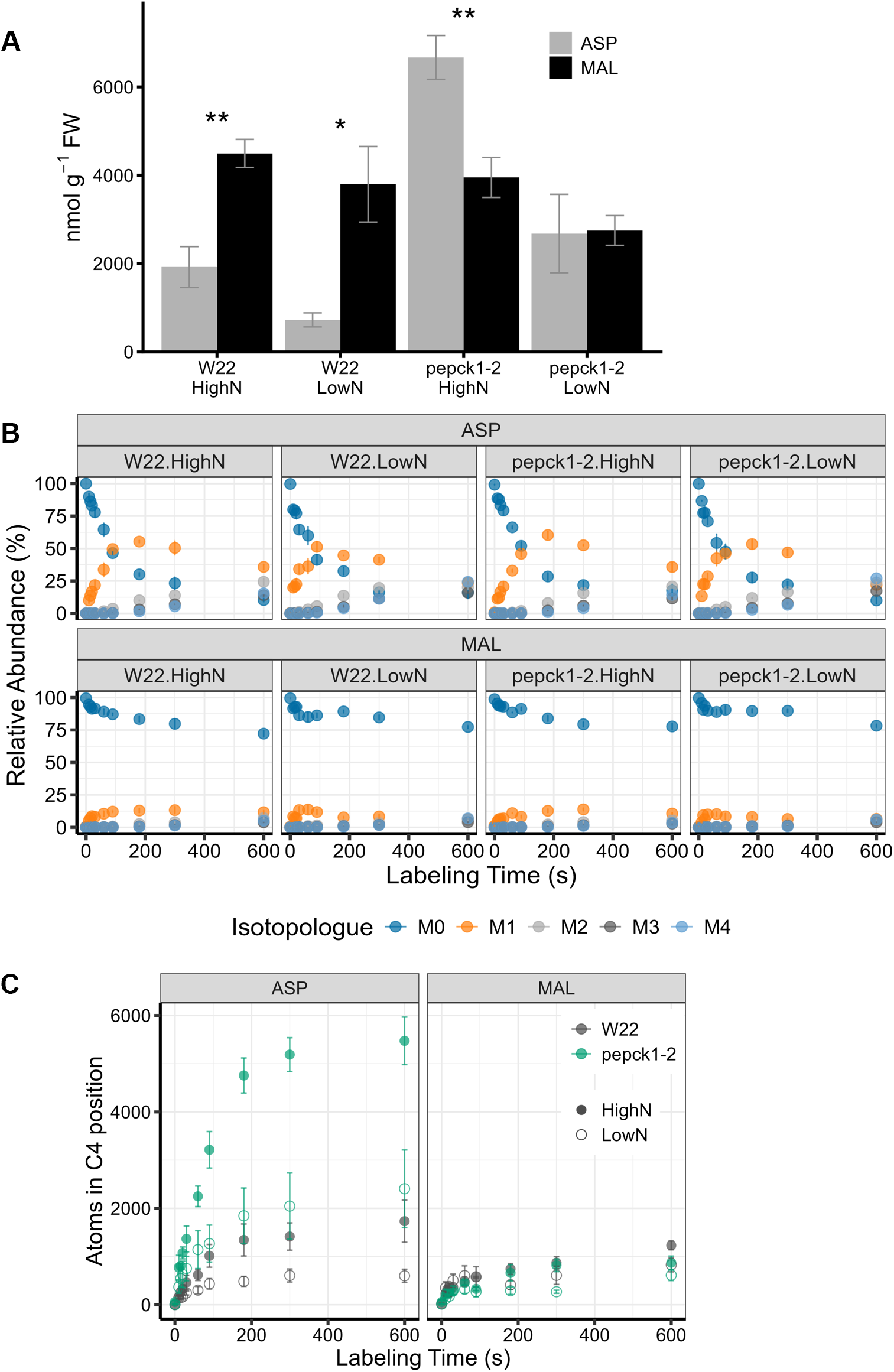
A.) Pool size of ASP and MAL in high and low N treatments in W22 and *pepck1-2* lines. B.) MID of ASP and MAL in high and low N treatments in W22 and *pepck1-2*. C.) Amounts of ^13^C in C-4 position of ASP and MAL, calculated using the method from Arrivault et al, 2017. Error bars represent means ± standard error. Significance assessed using two-sided Welch’s test. ***: p < 0.001, **: p < 0.01, *: p < 0.05. n = 4.

### Rapid isotopic labeling confirmed the *PEPCK* pathway was an active component in maize photosynthetic carbon assimilation

Stable isotope labeling was used to characterize differences in the metabolism of transfer acids in the NADP-ME and PEPCK pathways in greenhouse-grown plants at the V16 growth stage under replete fertilizer or low (no supplemental) nitrogen conditions. Synthetic air containing ^13^CO_2_ (400-410 ppm) with the balance of approximately 21% oxygen and 79% nitrogen was delivered to maize leaves by a custom-built small labeling chamber (∼40 mL volume) for various time intervals to label malate (NADP-ME pathway), aspartate (PEPCK pathway), and other intermediates involved in C_4_ and C_3_ photosynthesis. Malate and aspartate share a common precursor, oxaloacetate, and both serve as major C_4_ transfer acids. Their rapid initial labeling dynamics indicate flux through the NADP-ME and PEPCK pathways, respectively (Weissmann et al. 2016; AuBuchon-Elder et al. 2020). In both *pepck1-2* and W22, ^13^C was rapidly incorporated into malate and aspartate, with appreciable levels of singly labeled molecules within ten seconds, the shortest time point (Figure 3B), consistent with the expected fast rate of the initial steps of C_4_ photosynthesis (Weissmann et al. 2016; AuBuchon-Elder et al. 2020). At the 10, 20, and 30 second time points, both malate and aspartate were predominantly singly labeled, indicating ^13^C incorporation at the C-4 position (Figure 3B). Additional ^13^C label, e.g. in the C1-C3 positions, was delayed and incorporated into malate and aspartate at a slower rate (Figure 3B), reflecting conversions of C_3_ metabolism and Calvin-Benson Cycle (CBC) labeling of precursors (Suppl. Figure 3).

### Modeling of labeling trajectories and C4 calculation confirmed altered PEPCK metabolism in the mutant

Labeling was further analyzed using two approaches. First, malate and aspartate were compared using the M+1 isotopologue during the initial labeling period (0-30 s) to identify differences in early positional labeling in the fourth carbon (i.e., carbon-4). Then, the labeling pattern was scaled to metabolite pool size to assess potential differences in C_4_ metabolic activity between conditions for the labeling duration (Figure 3C) (Arrivault et al. 2017). While malate labeling showed little difference, aspartate labeling changed dramatically in the mutant and across nitrogen conditions, with labeling levels increasing most significantly in the high-nitrogen condition.

Second, ^13^C enrichment trajectories as calculated in (Kambhampati et al. 2024) were characterized for C_4_ and C_3_ metabolites using generalized additive models (GAMs) (Suppl. Figure 4) for the corresponding isotopologue distributions presented in Suppl. Figure 3. Labeling rates during the initial label incorporation (i.e. the first 30s) were compared among conditions using linear models, and pairwise differences in labeling slopes were assessed using Tukey-adjusted comparisons. Aspartate showed significantly greater early labeling rates in *pepck1-2* at high N than W22 at both high and low N, whereas no significant differences were observed for malate (Suppl. Figure 5).

### Grain biomass at maturity was increased in pepck1 plants grown in field conditions under both low and high N treatments

The effect of the PEPCK pathway mutation on productivity and the role of this pathway in coordinating N metabolism throughout the plant were evaluated in the field under limiting (no N fertilizer) or replete (150 kg Ha^-1^ N fertilizer) N supply. Biomass of stover, cob, and grain, as well as plant height were measured on mutant plants grown in Urbana, IL, during the summers of 2016-2022. Additionally, N concentrations were determined from a subsample of stalk and grain. Harvest index and NUE traits were calculated, and individual ears were used to assess yield component traits, including ear length, kernel number, and kernel weight (Suppl. Table 1A and 1B).

Stover biomass was significantly higher in *pepck1-2* than in W22 control plants across N treatments (p=0.026), and cob biomass was significantly higher in *pepck1-2* under high N (p=0.001) (Figure 4A). Mutant plants were also 6-9 cm taller than W22 (p=1.11E-8, Satterthwaite Type III ANOVA, Suppl. Table 1A), a phenotype also observed in greenhouse-grown *pepck1-2* plants at the V16 growth stage (Suppl. Figure 6). As anticipated, both vegetative and cob biomass were highly N-responsive across genotypes (p=2.66E-13 and 3.09E-12, respectively, Type III ANOVA with Satterthwaite’s method, Suppl. Table 1A).

**Figure 4:**
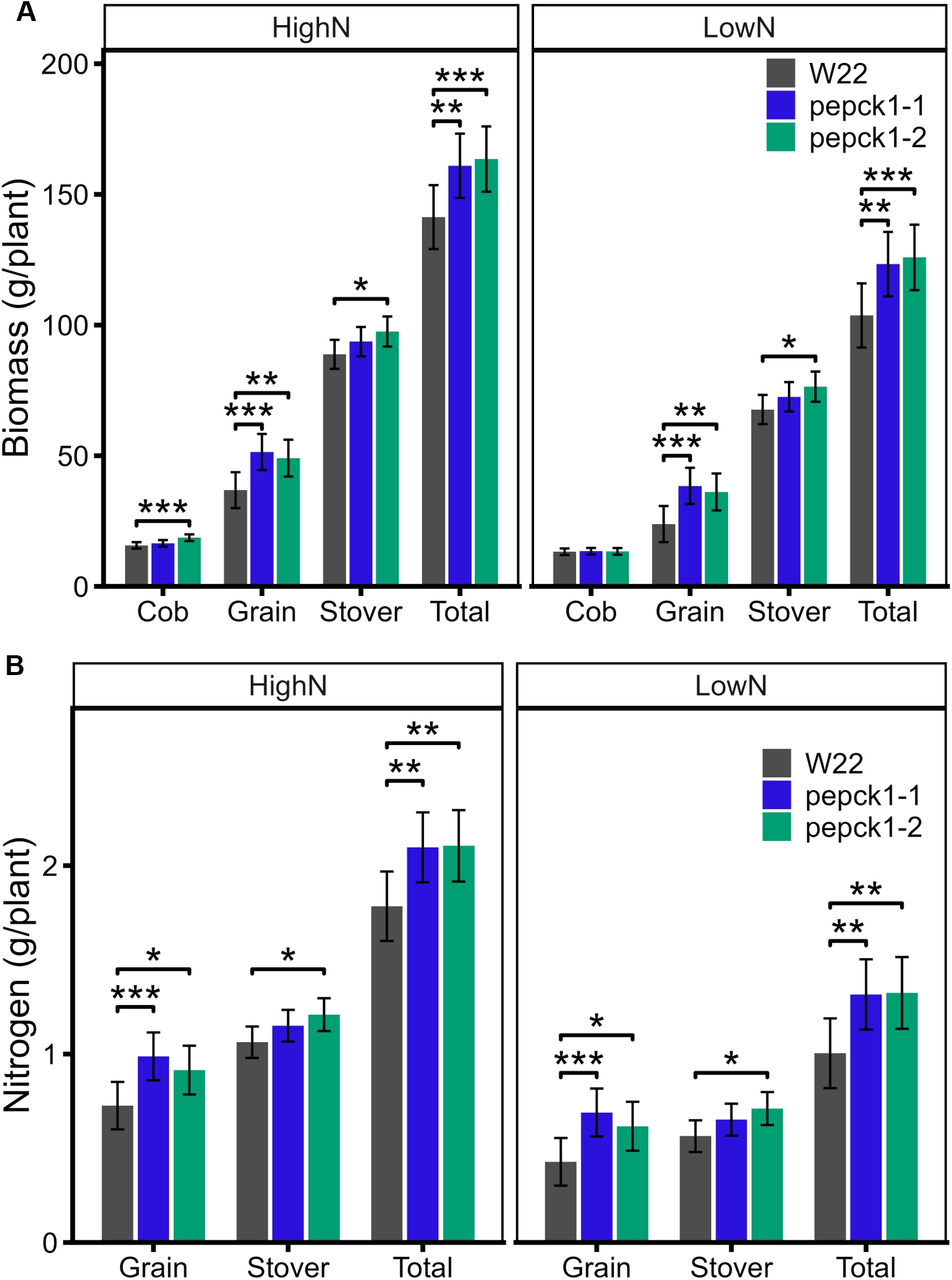
A.) Plant biomass and B.) nitrogen content at the R6 growth stage (physiological maturity) in the high and low nitrogen treatments. Data represent least-square means ± standard error for four years. For 2016 n = 2 plots, for 2017-2019 n > 4 plots per year, n= 3 plots in 2021 and 2022. Each plot is a measurement of 5 plants. ***: p < 0.001, **: p < 0.01, *: p < 0.05.

Grain biomass increased in both *pepck1* alleles relative to W22 under both high and low N (Figure 4A). An assessment of yield-component traits showed significantly longer cobs in the mutants; thus, larger and heavier cobs contribute to increased grain biomass (Suppl. Table 1B, Suppl. Figure 7). Consistent with their stronger impact on grain than stalk or cob biomass, harvest index (defined as grain biomass divided by total biomass) was significantly higher in the *pepck1* mutants (p=0.0003, Type III ANOVA with Satterthwaite’s method, Suppl. Table 1A).

At maturity, total N content was significantly higher in both *pepck1* mutants than in W22 under both high and low N (Figure 4B). Both mutants also accumulated significantly more N in the grain under each treatment, consistent with their greater grain biomass (Figure 4A, B). The increase in stover N content was specific to *pepck1-2*, which had significantly more stover N than W22 across N treatments (p=0.025). Neither *pepck1* mutant altered the nitrogen use efficiency of total biomass production, but the *pepck1-2* mutation significantly (p=0.003) increased nitrogen utilization (defined as the ratio of grain biomass to total plant N).

### Expression of nitrogen metabolism genes were increased in pepck1::Ds mutants

qPCR was performed to assess the impact of *pepck1::Ds* mutations on the expression of C_4_ pathway and known N metabolism genes in mature leaf segments from the V8 growth stage (Figure 5). Expression from *pepck1-1* and *pepck1-2* was compared to W22 controls using the delta-delta Ct method. Asparagine synthetase (AS3), which uses aspartate as a substrate, increased in expression in both *pepck1-1* and *pepck1-2*. Other genes closely related to aspartate metabolism (AS3, ASNase, AspAT-b) also increased by 1.2 to 2.2-fold in *pepck1-1*, but ASNase and AspAT expression did not change in *pepck1-2*. Other genes for enzymes in C_4_ photosynthesis showed slight decreases in expression (MDH3) or no change compared to W22 (PEPC, CA1, CA6).

**Figure 5:**
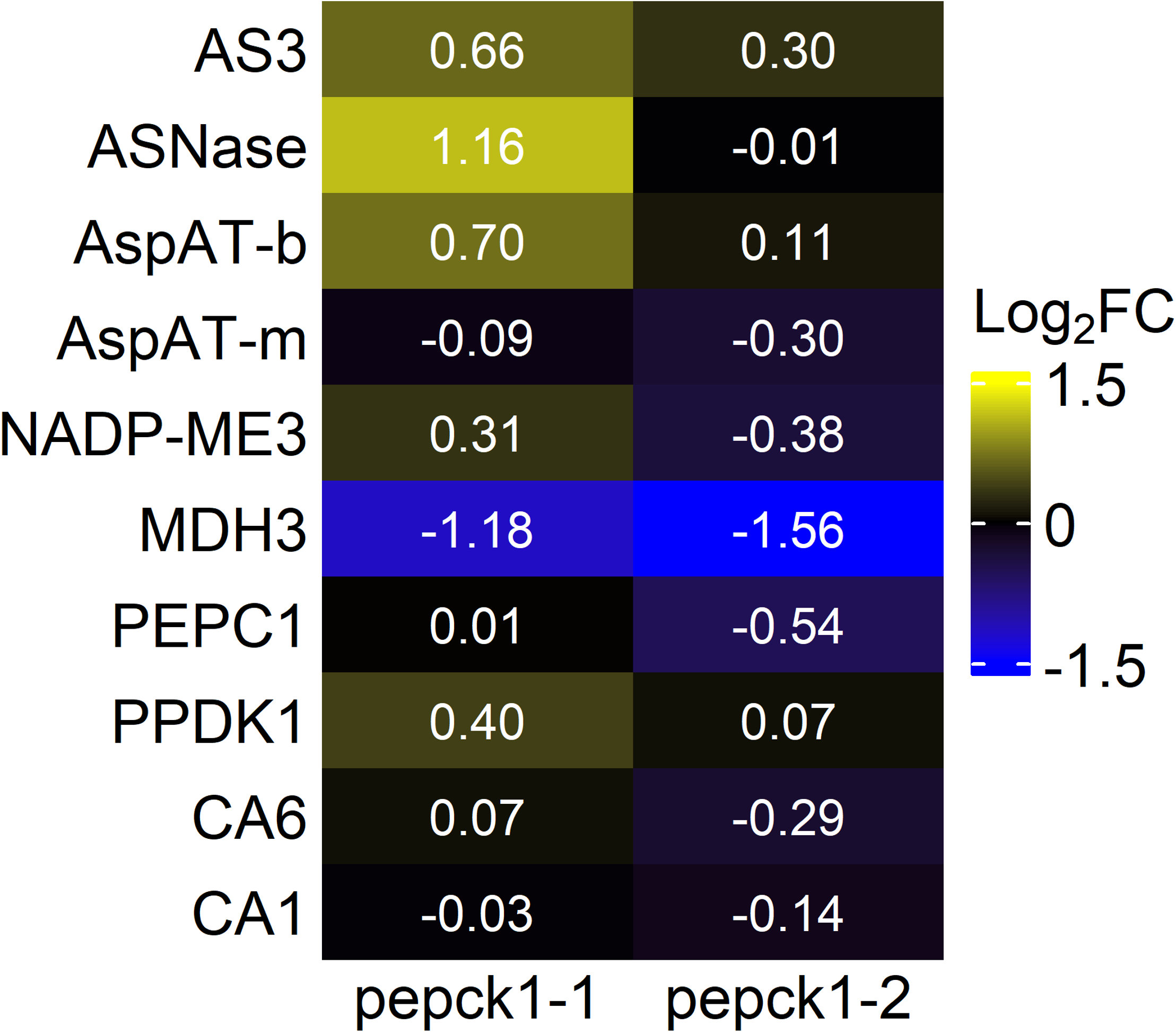
Expression of C_4_ pathway and nitrogen metabolism genes in *pepck1-1* and *pepck1-2* mutants compared to W22 in mature leaf at V8 using qPCR. Asparagine synthetase3 (Zm00004b001218, AS3), L-asparaginase (Zm00004b006101, ASNase), aspartate aminotransferase bundle sheath isoform (Zm00004b018905, AspAT-b), aspartate aminotransferase mesophyll isoform (Zm00004b015382, AspAT-m), NADP-malic enzyme (Zm00004b015828, NADP-ME3), malate dehydrogenase3 (Zm00004b019483, MDH3), PEP carboxylase (Zm00004b032576, PEPC1), pyruvate, orthophosphate dikinase (Zm00004b030561,PPDK1), carbonic anhydrase6 (Zm00004b027167, CA.6), carbonic anhydrase1 (Zm00004b019533, CA1).

### Photosynthetic traits were unaffected in the pepck1::Ds mutants

Net CO_2_ assimilation (*A*_net_) was measured using a photosynthetic gas exchange instrument (Li-6800, LI-COR Inc, Lincoln NE) on the top expanded leaf of plants approximately 30 days after planting in the greenhouse, approximately ¼ of the way down the leaf length from the tip. The initial slope of the CO_2_ response curve decreased slightly in *pepck1-1* compared with W22. The maximum *A*_net_ in *pepck1-1* trended lower but was not statistically different from W22 (Figure 6A).

**Figure 6:**
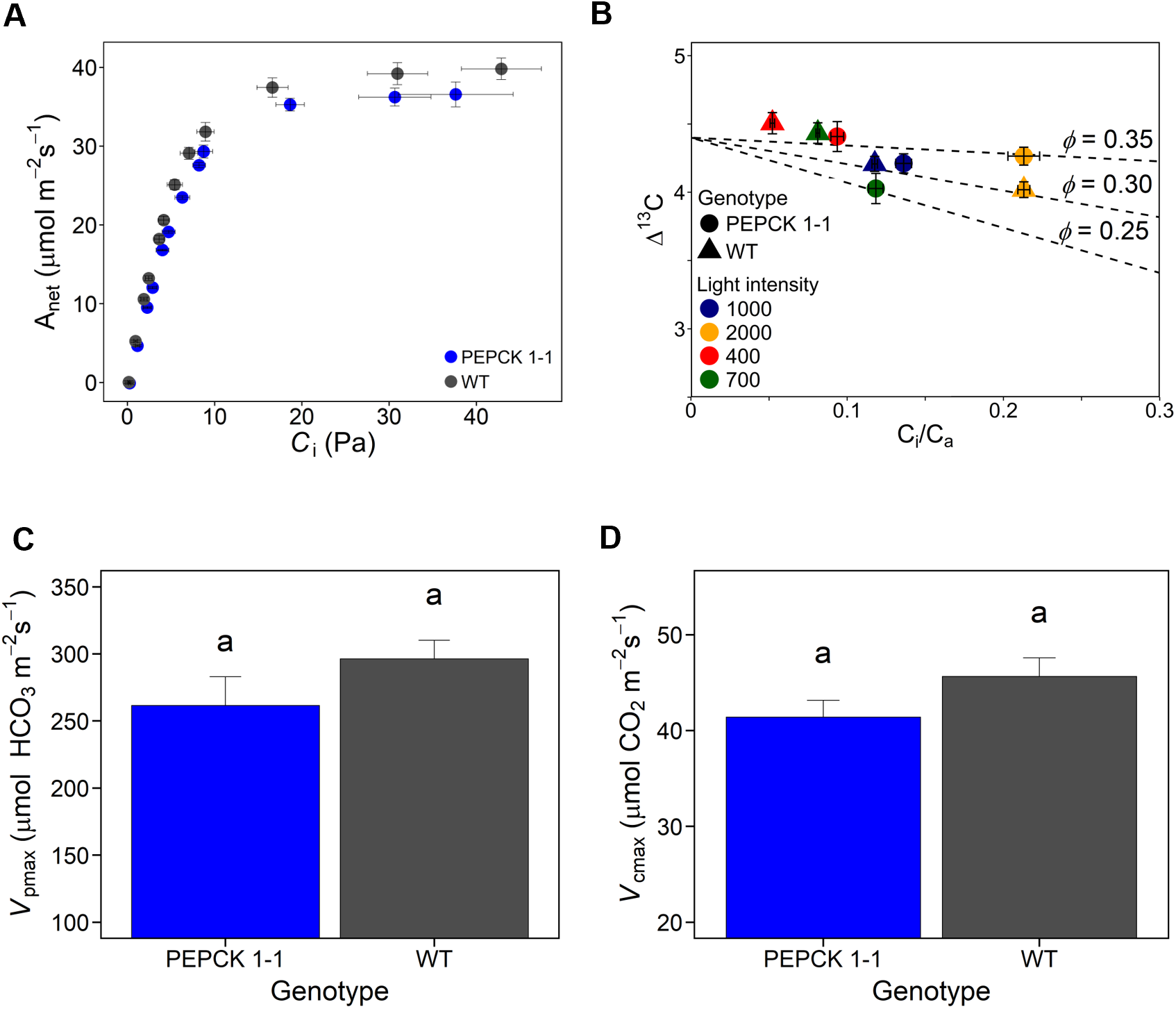
Gas exchange and enzyme activity in *pepck1-1* and W22 control plants. A.) Plants were grown for 30 days in Pullman, WA, and A_net_-C_i_ curves measuring the response of A_net_ and C_i_ to a gradient in ambient CO_2_ concentration. B.) ^13^C discrimination in *pepck1-1* and W22 plants was measured under a gradient of stable light conditions (400, 700, 1000, 2000 μmol photons m^-2^ s^-1^). Dashed lines represent theoretical relationship between Δ^13^C and *C*_i_/*C*_a_ for leakiness values: 0.25, 0.30, 0.35. C-D.) Enzyme activity for phosphoenolpyruvate carboxylase (PEPC) and rubisco were measured in *pepck1-1* and W22.

With decreased flux through the PEPCK pathway, we hypothesized that the bundle sheath CO_2_ concentration and Calvin-Benson Cycle (CBC) activity would be altered (Bellasio and Griffiths 2014; Kromdijk et al. 2014). However, *pepck1-1* and W22 did not differ in the relationship between carbon discrimination (Δ^13^C) and *C*_i_/*C*_a_ across stable light intensities (400, 700, 1000, and 2000 μmol photons m^-2^ s^-1^), indicating that an approximate 50% reduction of the PEPCK pathway in the *pepck1-1* mutant did not influence bundle sheath leakiness over the observed range of *C*_i_/*C*_a_ (Figure 6B). Measurements of *C*_i_/*C*_a_ were generally low but within the range observed for maize genotypes and conditions, which can vary widely and depend on the methodology (von Caemmerer et al. 2014; Kolbe and Cousins 2018; Crawford et al. 2024). Here, relatively low growth light and temperature conditions would acclimate to a low *g_s_* (and *C*_i_/*C*_a_). In addition, measurements were based on maintaining a constant CO_2_ concentration in the sample, whereas some reports control the CO_2_ concentration in the reference, which can result in reduced *C_a_*values due to CO_2_ drawdown and in inflated *C*_i_/*C*_a_ ratios. In the mutant, carboxylation reactions (PEPC and Rubisco) decreased modestly, which was qualitatively consistent with changes in *A*_net_ (Figure 6C-D). Further studies of net assimilation under varying light conditions were attempted repeatedly as presented in the supplemental material but were inconclusive, possibly due to differences in seasonal conditions at the time of measurement or variability in leaves (Suppl. Figure 8). The absence of changes in photosynthetic rates in mutants suggests that the photosynthetic role of PEPCK is small relative to NADP-ME in maize for the conditions tested here.

### PEPCK activity in developing kernels was not influenced by the mutations

Because photosynthetic measurements could not explain differences between W22 and the mutants observed in the field, we measured PEPCK activity in developing kernels. Although *pepck1* is implicated in C_4_ leaf photosynthesis (Yi et al. 2019; Yang et al. 2026), a small pleiotropic effect in the kernel could affect the sink capacity of the ear and thus potentially explain the observed changes in yield. As C_4_ photosynthetic enzymes operate with considerably higher kinetic activity than the anaplerotic C_3_ isoforms, the presence of PEPCK activity in the seed, including a small contribution from PEPCK1, could be counterproductive to the import of seed assimilates and negatively impact grain yield. Analysis of seed-based activity showed there was not a significant difference between pepck1 and W22 at any of the three developmental time points (8, 16 and 32 days) post-pollination (p = 0.738, 0.127, and 0.472, Figure 7), although in every case the average measured value was less than W22.

**Figure 7:**
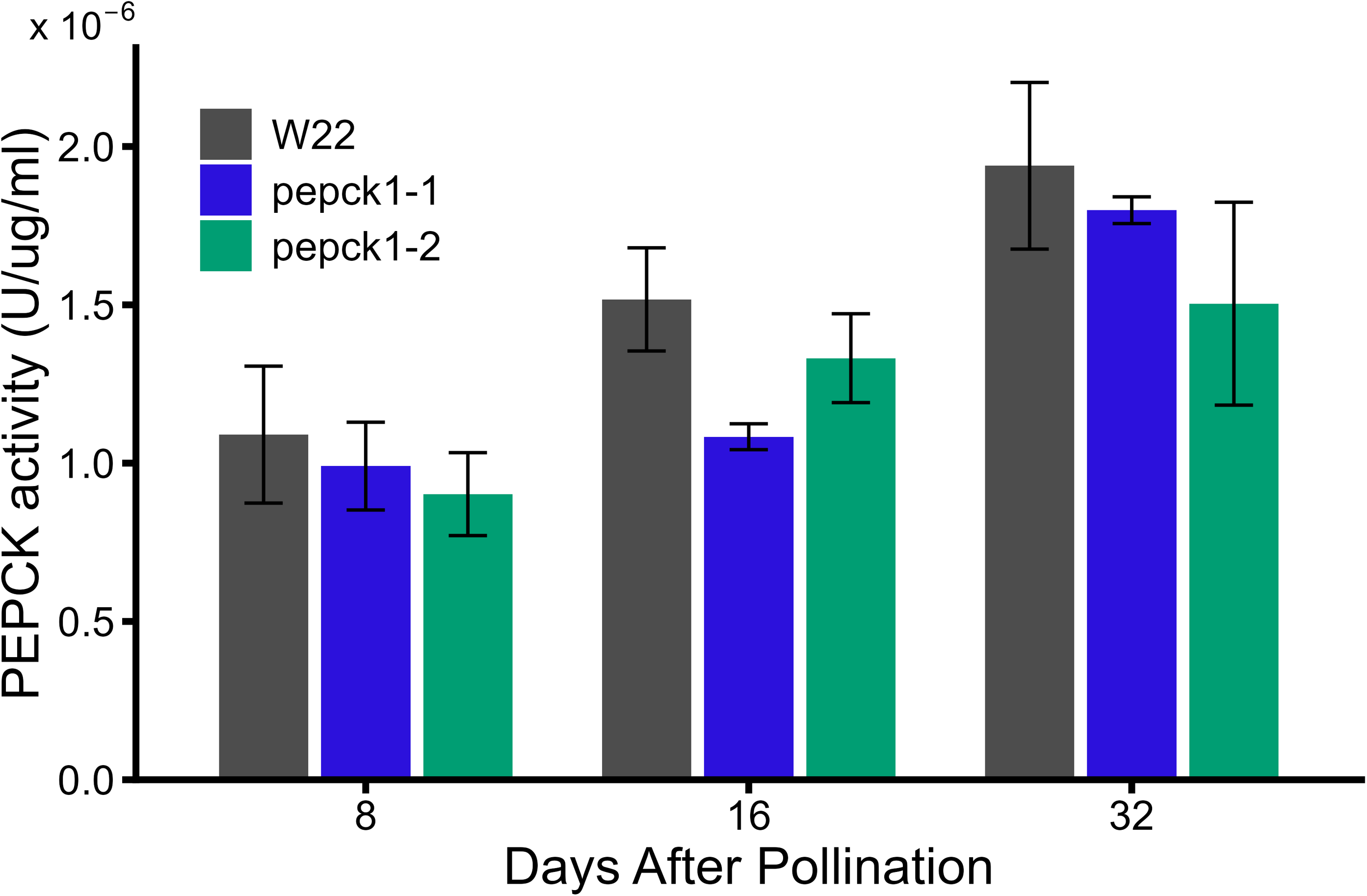
PEPCK enzyme activity in *pepck1-1* and *pepck1-2* in seeds. Error bars represent means ± standard error. Compact letter assignment represents significant differences (p < 0.05) among the three genotypes. No letter is displayed if there is no difference. (n = 3, ANOVA followed by Tukey’s multiple comparisons test).

## Discussion

Nitrogen fertilizer, together with improved genetics and agronomic practices, has significantly increased US crop yields over the past 75 years. However, N fertilizer is typically applied in excess of plant demand, which has undesirable side effects beyond added costs, such as eutrophication of waterways and increased greenhouse gas emissions (Bijay-Singh and Craswell 2021; de Vries 2021; Paudel 2025). Improving maize yield and NUE while reducing N inputs would enhance US agriculture’s profitability, benefit the environment, and increase yields for small-holder farmers in other countries. N metabolism is intimately linked with photosynthetic carbon assimilation in C_4_ crops, which rely on species-dependent combinations of C_4_ sub-types, of which the NADP-ME subtype is considered the most N-efficient (Ghannoum et al. 2005; Pinto et al. 2014, 2016). Maize is thought to contain both NADP-ME and PEPCK pathways; thus, decreasing the latter could enhance productivity by reducing the need to synthesize additional abundant photosynthetic enzymes to operate both pathways, or by reducing the energetic costs of cyclic nitrogen shuttling, resulting in improved NUE. It remains unclear whether the two pathways complement each other under changing environments (Furbank 2011; Stitt and Zhu 2014) but photosynthetic performance in mutant lines was unchanged from W22 under the limited conditions explored here. Until now, no genetic studies have confirmed two active C_4_ photosynthetic pathways in maize, leaving questions about nitrogen use and photosynthetic contribution unresolved. To understand the role of the PEPCK pathway relative to the NADP-ME pathway in maize, we generated transposon mutants of the higher expressed *Pepck* gene (*Pepck1*). Two *pepck1::Ds* mutants were confirmed by DNA sequencing, RNA expression, and enzymatic assays of PEPCK to be knockdowns. The *pepck1*::*Ds* mutants showed decreased *Pepck1* gene expression and enzyme activity compared to W22 controls, with *pepck1-2* having a stronger effect than *pepck1-1* (Figure 2). NADP-ME did not increase in expression, and other enzymes in the NADP-ME pathway did not increase gene expression to compensate for decreased PEPCK activity (Figure 5).

The current study with *Pepck1* transposon mutants shows that PEPCK1 catalyzes a decarboxylation of carbon shuttled as aspartate, as an alternative route to decarboxylation by NADP-ME in the nominally NADP-ME subtype C_4_ species maize, providing genetic support for two-pathway operation. Changes in aspartate and alanine pool sizes and metabolism in these *Pepck1* transposon mutants were consistent with disruption in the C_4_ metabolic pathway but did not affect photosynthetic traits. This may reflect a largely redundant role for PEPCK in photosynthesis, at least under the conditions explored here, where leaf PEPCK activity was reduced by approximately 50-75% throughout plant development (Figure 2). In contrast, multiple years of field trials in low-N and high-N conditions showed increased seed yield in mutants, consistent with improved NUE and a role for PEPCK1 in coordinating these two pathways, and indicating the potential importance of this pathway for both sustainable and subsistence farming efforts.

The accumulation of aspartate in the leaves of *pepck1*::*Ds* mutant plants was the most striking metabolic change and is consistent with decreased PEPCK decarboxylation (Figure 1, Figure 3A). ^13^C was rapidly incorporated into aspartate because aspartate generation occurs upstream of the pathway mutation (Figure 3B) and the aminotransferase is subject to significant exchange. Importantly, the aspartate change, resulting from PEPCK alteration, provides genetic evidence that PEPCK, not NAD-ME, is the predominant secondary pathway that shuttles aspartate in maize, consistent with prior descriptions based on biochemical evaluations (Furumoto et al. 1999; Wingler et al. 1999; Majeran et al. 2010; Pick et al. 2011). The aspartate pool size was N-responsive in both the mutants and W22; however, the mutant aspartate pool at low N was already larger than the aspartate pool in W22 at either N condition (Figure 3A). Both the aspartate pool size and labeling rates were increased dramatically for the *pepck1-2* mutant grown with higher N supply, indicating a greater role for PEPCK decarboxylation under higher N supply. *Pepck1* mRNA expression is not highly N-responsive in leaves from diverse maize inbreds or hybrids (Cheng et al. 2021) suggesting PEPCK1 enzyme activity likely modulates aspartate responses to N. Flux through PEPCK1 may be more important when N supply is high, and leaves increase not only photosynthesis but also their rate of nitrate assimilation. The further elevation of aspartate levels in the *pepck1-2* mutant grown at high N is associated with stronger increases in vegetative and grain biomass, and eventually nitrogen utilization.

Although the role of accumulated aspartate in the mutant is unknown, aspartate could serve as a substrate for many other non-C_4_ photosynthetic cellular roles, notably N metabolism and protein synthesis (Galili 2011) or as a signaling molecule conveying the leaf’s N status to the rest of the plant (Seebauer et al. 2004). Aspartate has previously been shown to preferentially participate in roles outside the C_4_ shuttle under N-limiting conditions (Khamis et al. 1992). In our experiments, both total malate concentrations and malate involved in the C_4_ shuttle (Figure 1) were stable across N treatments and between the *pepck1-2* mutant and W22 control, indicating that aspartate is the more N-responsive carbon shuttle in C_4_ photosynthesis, at least under these conditions and genotypes.

It was intriguing to consider whether a small amount of PEPCK1 activity could contribute to active seed assimilation metabolism. Average seed activity was lower in mutants than in W22; however, the difference was not statistically significant. It remains unclear how a larger aspartate pool specifically contributes to productivity, but investigations into other PEPCK pathway mutations are currently underway to better understand the pathway’s complexity.

## Conclusion

Here, we provide genetic evidence that the PEPCK pathway is active in C_4_ photosynthesis, using transposon mutant analysis, including isotopic labeling and pool-size differences that are specifically and temporally linked to the rapid kinetics of the C_4_ pathway. Evidence accumulated over multiple field seasons demonstrated that modifications to PEPCK pathways can increase yield, promoting stronger grain biomass responses to N fertilizer, without changing carbon assimilation. Possible benefits to the plant of using the PEPCK pathway, and why it has been maintained throughout evolution and across decades of breeding, remain open questions. In less ideal growing conditions, such as subsistence farming or when fertilizer is not applied, these results are promising for altering nitrogen use and improving grain yield.

## Methods

### Mutant generation and Ds reverse genetic screening

Mutagenesis of the *Pepck1* locus in maize (Zm00004b001002) was performed using the *Ac/Ds* tagging stock B.S07.0675R, located 601.7kb 3’ of the gene, as the donor *Ds* element. Because of the large distance between the donor *Ds* element and the *Pepck1* locus, the *Ds* element was remobilized in a two-step process: first to a location closer to *Pepck1*, then to a second population. In the first round, we screened a population of 3,150 individuals from a heterozygous pollen donor (1,575 chromosomes). Populations for *Ds* screening were generated by crossing to W22::*r-sc:m3/r-sc:m3*, and *Ds* element transposition into *Pepck1* was performed as previously described (Kolbe et al. 2018). A single heritable insertion allele was found in a gene (Zm00004b001005) ∼140kb from *Pepck1*. In the second round, we screened a population of 800 individuals from a heterozygous pollen donor, with the new insertion allele serving as the donor *Ds* element. Two heritable insertion alleles were identified in *Pepck1* and confirmed by Sanger sequencing of the PCR-amplified gene using a *Ds*-specific primer. Mutants were confirmed in subsequent generations using a gene-specific primer (CGTCCACGAGCATAGACCTC) and a *Ds*-specific primer (GTTCGAAATCGATCGGGATA) to identify plants carrying the mutant allele. A second gene-specific primer (TTACACAGCGTCATGTCTCAACT) downstream of the *Ds* insertion was used in combination with the first gene-specific primer to identify plants lacking the *Ds* insertion.

### Mutant PEPCK enzyme activity assay over plant development

Maize seeds of W22, *pepck1-1*, and *pepck1-2* lines were germinated in plug trays (jolly gardener:turface 3:1 soil blend) and later transferred to a standard potting mix (Berger BM7-35%) after 14 days. The greenhouse was maintained under standard maize conditions for growth at day length 16 hours at approximately 28°C Day/22°C night, 40-50% humidity, with supplemental light as needed. To determine PEPCK activity, nine developmental time points were sampled from six biological replicates (i.e., leaves from six independent plants at each time point, n=6) for each line. The top leaf of the canopy was sampled, avoiding the midrib approximately ¼ of the way from the tip of the leaf (similar to the location for photosynthetic and labeling studies), and rapidly quenched using liquid nitrogen while being careful not to shade the leaf and stored at -80°C. The subtending leaf (leaf below the developing ear) was similarly sampled at 8, 16, and 32 days after pollination. Samples were finely ground in liquid nitrogen with a mortar and pestle.

The PEPCK activity was measured using the method outlined by (Sharwood et al. 2014, 2016). Approximately 100mg of frozen, ground tissue was taken for extraction. 1mL of cold extraction buffer containing 50mM HEPES of pH of 7.0, 5mM freshly prepared DTT, 1% (w/v) PVPP, 2mM EDTA, 2mM MnCl_2_, and 0.05% Triton was added to the samples. The sample was then vortexed for 30s and subsequently homogenized for 2min using a homogenizer to ensure proper extraction. Samples were then left on ice for 10min. Next, the crude extract was centrifuged at 13000g, 4°C for 30s. PEPCK assay buffers were prepared fresh as 50mM HEPES, pH 7.0, 4% mercaptoethanol (w/v), 100mM KCl, 90mM NaHCO_3_,1mM ADP, 2mM MnCl_2_, 0.14mM NADH, 15mM PEP and 6 units of malate dehydrogenase. MgCl_2_ was absent from the extraction buffer and the PEPCK reaction buffer to avoid interference from PEPC and other enzymes (Sharwood et al. 2016). Commercial standard PEP (phosphoenolpyruvate) concentrations ranging from 5mM to 20mM were used to validate and optimize the PEPCK assay. PEPCK activity was measured at 340nm using a Tecan i-control spectrophotometer. Samples were normalized by measuring total extracted protein using the Bradford assay with external standard curves based on commercial bovine serum albumin.

### qPCR analysis

First-strand cDNA synthesis was performed using 1µg RNA using igScript Reverse Transcriptase from Intact Genomics (Saint Louis, MO; Cat #3333) using random hexamers. Genes related to C4 photosynthesis and aspartate-asparagine cycling were quantified; the primers are listed in Suppl. Table 2. qPCR was performed using ig SYBR Green qPCR 2x Master Mix from Intact Genomics (Saint Louis, MO; Cat #3356) on a Roche Light Cycler 480 Real-Time PCR system. Differential gene expression was measured by comparing C4 pathway genes with the expression of *NAC* and *UPF1* housekeeping genes for maize (Lin et al. 2014). Relative quantification of *Pepck1* gene expression in both *pepck1* mutants was compared to the W22 control using the delta delta Ct method.

### Gas exchange and photosynthetic measurements

Plants used for gas exchange measurements at Washington State University (Pullman, WA) were grown in a greenhouse during October and November. Growth conditions included a minimum of 600 µmol m^-2^ sec^-1^ light, mean day and night temperatures (± standard deviation) of 25.6 ± 1.3°C and 22 ± 1.5°C, respectively, and a mean relative humidity (RH) of 44.9 ± 11.2%. Seeds were planted in 11.3L pots filled with potting soil (Sunshine LR1 mix, SunGro, Agawam, MA, USA). Plants were fertilized with 2.8g/L 20-20-20 + micronutrients (JR Peters Inc.) every other watering and with 1.3 g/L iron chelate (Sprint 330, BASF) at 14 and 20 days. *A*_net_-*C*_i_ curves were quantified at a leaf temperature of 28°C, 72-75% RH, and 2000µmol m ² s ¹ light. The youngest, fully expanded leaf was placed in a 3cm by 3cm leaf chamber of the LiCor Li-6800 while attached to the plant. CO_2_ concentration in the leaf chamber varied as follows: 400, 0, 50, 100, 125, 175, 200, 250, 300, 500, 750, 1000 µmol mol^-1^ air.

For carbon discrimination (Δ^13^C) measurements, four plants of each W22 and *pepck1* genotypes were measured with a LI-COR Li-6800 gas exchange instrument, and the exiting air was analyzed by a tunable diode laser (TDL-AS; Campbell Scientific, Logan, UT, USA). The gas exchange chamber was maintained at 31°C, and the CO_2_ concentration in the sample was 400µmol mol□¹. Gas exchange and Δ^13^C were measured for each plant at photosynthetically active radiation (PAR) of 2000, 1000, 700, and 400 µmol quanta m□² s□¹. The leaf remained at each light intensity until steady state was reached, and 10 cycles were recorded at 2min 20s intervals. In each cycle, the TDL-AS measured carbon isotopes in both reference (inlet) and samples (outlet) air from the gas exchange chamber. Two calibration tanks (Liquid Technology) were used to calibrate for gain and possible offset in the TDL-AS measurements [see (Bowling et al. 2003; Ubierna et al. 2013)]. Photosynthetic ^13^C discrimination was calculated as (Evans et al. 1986):

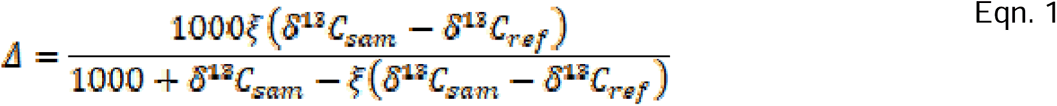

Where δ^13^C_sample_ and δ^13^C_ref_ are the carbon isotope composition of the leaf sample (chamber) and reference air of the Li-6800, respectively. The ξ values were calculated as ξ = *C*_ref_ /(*C*_ref–_-*C*_sam_), where *C*_ref_ and *C*_sam_ are the CO_2_ concentrations of dry air entering and exiting the leaf chamber, respectively. The delta notation expressed in ‰ was relative to the Vienna Pee Dee Belemnite carbon standard (Evans et al. 1986).

*In vitro* phosphoenolpyruvate carboxylase (PEPC) and ribulose-1,5-bisphosphate carboxylase-oxygenase (Rubisco) activities were measured at 25°C in leaf tissue frozen in liquid N_2_ immediately after sampling. Leaf tissue was ground uniformly for 2 min using a pre-chilled mortar and pestle in 1mL of 100mM HEPES, (pH 7.8), 10 mM dithiothreitol, 1% polyvinylpolypyrrolidone, 1mM EDTA, 0.1% (v/v) Triton, and 2% (v/v) of a plant protease inhibitor cocktail (P9599; Sigma-Aldrich, St Louis, MO, USA). A pinch of sand was added to facilitate grinding. The extract was centrifuged for 10s at 4°C, and the supernatant was collected. Enzyme activity assays were initiated immediately. Rubisco and PEPC activities were measured at 30°C using spectrophotometric assays that monitored NADPH consumption at 340nm, as described by Sharwood and Studer (Studer et al. 2014; Sharwood et al. 2016).

Two fluctuating light experiments were conducted on one-month-old plants. Eight biological (whole-plant) replicates in the greenhouse in Pullman, WA were assessed simultaneously with the other photosynthetic measurements described above. Gas exchange measurements were made on the youngest, full-expanded leaf of six W22 and seven *pepck1-1* plants ¼ of the way down the leaf from the tip. Two Li-6800 gas exchange instruments were used to measure the effect of fluctuating light conditions on net photosynthetic rate (*A*_net_). In the Li-6800 chamber, leaf chamber conditions included a temperature of 31°C and a sample CO_2_ concentration of 450 µmol mol ¹. While stomatal conductance and *A*_net_ were stabilizing, light intensity remained at 1500 µmol quanta m□² s□¹. Once stable, light intensity alternated between 1500 and 250 µmol quanta m□² s□¹ at the following intervals: 10 min at 1500, 10 min at 250, 10 min at 1500, then followed by 14 1-minute intervals of alternating 250 and 1500 light intensities. *A*_net_ was measured every second throughout the experiment. This experiment was repeated at a different location and in a different environment to mimic the longer days of the Midwest growing season using *pepck1-1* and *pepck1-2* alleles at the Donald Danforth Plant Science Center Plant Growth Facility (St. Louis, MO) in April-June 2021, with final measurements made in June 2021. Plant growth conditions in June were > 25°C during the day and 22°C at night, reflecting summer in St. Louis, MO, ∼50% relative humidity, and a minimum of 500 µmol m^-2^ s^-1^ light.

### ^13^CO_2_ labeling

Plants were grown in the greenhouse at the Donald Danforth Plant Science Center in Berger BM7-35% bark medium in 2.5-gallon pots under normal fertilization or no fertilization, from December 15, 2018, to February 6, 2019. Plants were grown at 28°C during the day and 22°C at night in greenhouses, with a minimum relative humidity of 40%, and 14-hour days, 10-hour nights, and supplemental light provided when sunlight was below 600 µmol m ² s ¹. Plants were sampled 55 days after sowing.

^13^CO_2_ was delivered to the leaf in a premixed gas at atmospheric concentrations of O_2_ and N_2_ and at 400ppm ^13^CO_2_. The flow rate was 0.5mL/min, and the gas was supplied to the underside of the maize leaf within an approximately 40mL labeling chamber, with an outgassing hole on the top side of the chamber. Plants were sampled at 0, 10, 15, 20, 30, 60, 90, 180, 300, and 600s. Leaf samples approximately ¼ of the way down from the leaf tip were snap-frozen with a freeze clamp, submerged in liquid N_2_, midribs removed, and kept at -80°C until further processing.

### Metabolite extraction

Metabolite dynamic labeling, pool size measurement, and extracellular metabolite data were collected and confirmed on two LC-MS systems using the method of (Czajka et al. 2020). All samples were extracted with a 7:3 (v:v) methanol:chloroform solution containing acid-washed glass beads at 4°C for 6 hr, with the solution vortexed hourly, 4 parts ddH_2_O were added (7:3:4, v/v), and the upper aqueous phase was collected and centrifuged through 3kDa filters at 0°C. Samples were then frozen, lyophilized, and reconstituted in a 1:1 (v:v) methanol:ddH_2_O solution. We added 5μM PIPES as an internal standard before extraction and used it to normalize samples.

### Chromatography and mass spectrometry conditions

A Shimadzu Prominence-xR UFLC system was used for chromatographic separation, and a SCIEX hybrid triple quadrupole-linear ion trap MS equipped with a Turbo V^TM^ electrospray ionization (ESI) source detected metabolites using the method described by (Czajka et al. 2020; Kambhampati et al. 2021). The mobile phase solvents, A and B, contained 10mM ammonium acetate and 5μM medronic acid in water and 9:1 acetonitrile:ddH_2_O, respectively. A 3μL sample was injected onto an InfinityLab Poroshell 120 HILIC-Z (2.1 x 100 mm, 2.7 µm, Agilent Technologies) column held at 40°C, and the following gradient was used: the initial concentration of 95% B was linearly decreased to 70% B over 8 min, then to 50% B over the next 4 min. The gradient was then ramped to 30% B over 0.5 min, held for an additional 1.0 min for cleanup, and then ramped back to 95% B over 0.5 min. A 6 min equilibration was used to return the column to starting conditions before the next injection. The total runtime was 20 min, and a flow rate of 250μL/min was used throughout. A polarity-switching method was used to detect sugar phosphates, sugars, organic acids, and mevalonate pathway intermediates in negative ionization mode, and amino acids in positive ionization mode, using a targeted MRM approach. The source conditions used were as follows: ion spray voltage, 4.5 kV (ESI+ and ESI-); ion source temperature, 400°C; source gas 1, 45; source gas 2, 40; and curtain gas, 35. Data for absolute quantifications were analyzed using the quantitation wizard available in Analyst (v. 1.6.2) software (SCIEX, Concord, Canada). ^13^C enrichment measurements were quantified by manual peak integration. Measurements were corrected for the natural abundance of ^13^C using IsoCorrectoR (Heinrich et al. 2018).

### Normalization of malate labeling to account for the inactive pool

Malate and aspartate pools differed in size and in their participation in the PEPCK and NADP-ME pathways, which had to be accounted for to assess C_4_ metabolism. As in other reports (Weissmann et al. 2016; Arrivault et al. 2017; AuBuchon-Elder et al. 2020), malate labeling in this experiment was incomplete, indicating a large inactive malate pool in the maize leaf that did not participate in photosynthesis. After 10 min of ^13^C labeling, over 70% of the malate pool remained completely unlabeled at all carbon positions across all four genotypes and N treatments (Suppl. Figure 3). No other measured metabolite showed a large inactive pool; for example, 90% of the aspartate pool became at least partially labeled within the same time. We calculated the active malate pool by correcting for the unlabeled portion at the longest time point for each of the four genotype-by-N treatments (i.e., *pepck1-2* at either high or low N and W22 at either high or low N). The unlabeled fraction, M_0_, from a label-saturated time point was subtracted from malate M_0_ isotopologues for each time point, and the remaining M_0_ was then rescaled with other isotopologues to estimate the active C_4_ malate pool. Ten minutes of labeling with ^13^CO_2_ was sufficient to approximate the active malate pools in maize leaves, as labeling levels did not change appreciably from 10 to 60 min (Arrivault et al. 2017).

### Evaluation of N-responsive phenotypes in field experiments

Field experiments were conducted at the Department of Crop Sciences Research and Education Center in Champaign, IL during the 2016, 2017, 2018, 2019, 2021, and 2022 growing seasons on plots shown to be responsive to N (Uribelarrea et al. 2009; Cheng et al. 2021). The soil type was a Drummer silty clay loam, with a pH of 6.2. Maize W22 plants were grown in a split-plot design with individuals in each main plot (6 rows, 5.33m long, 76 cm row spacing) paired in adjacent rows that received either 150 kg Ha^-1^ fertilizer N (replete N) or no exogenous N (low N). Based on N recovery from plots with no applied fertilizer at this field site, the soil N capacity was 75kg ha^-1^, and the site was subjected to a corn-soybean rotation to homogenize soil N. Fertilizer was applied as granular ammonium sulfate in a narrow band adjacent to the root zone at the V3-V4 plant growth stage. Plots were maintained weed-free by a pre-plant application of herbicide (atrazine + metolachlor), followed by hand cultivation as needed. Plants were hand-pollinated to ensure adequate fertilization.

Biomass sampling was conducted at physiological maturity (R6 growth stage) using a protocol modified from (Haegele et al. 2013). Above-ground biomass from five representative plants was collected from the center of each plot and separated into ear (grain and cob) and stover (leaf, stem, and husk) fractions. Before chipping, two plants from the stover fraction were oriented in the opposite direction to the remaining three to improve the homogeneity of the chipped plant sample during the chipping process. The fresh weight of the stover fraction was measured, and the sample was shredded in a commercial brush chipper (Vermeer BC600XL; Vermeer Corporation). A representative 100-250g aliquot of the chipped material was weighed and dried to calculate a moisture adjustment and determine the dry weight of the total stover sample. The stover aliquot was further ground using a Wiley mill (Thomas Scientific) to pass through a 2mm mesh screen, then analyzed for percent N by combustion analysis in a Fisons NA 2000 N Analyzer. Ear samples were dried to 10% moisture and hand-shelled to partition and weigh the grain and cob. Whole kernel samples were analyzed on a Perten DA 7200 Near Infrared (NIR) analyzer to determine moisture, starch, oil, and protein concentrations (Ramchandran et al. 2017). Grain N was calculated from percent protein using a conversion factor of 6.25, as used in the Kjeldahl method (Jones 2018). Total N utilization was calculated as total biomass divided by total plant nitrogen. Grain N utilization was calculated as grain biomass divided by total plant N. Grain yields were mathematically adjusted to 0% moisture.

### Statistical Analysis

Differences in phenotypic and yield component traits (Suppl. Tables 1A and 1B) were analyzed using linear mixed-effects models implemented in R using the lmer and lmerTest packages (Bates et al. 2015; Kuznetsova et al. 2017). Genotype and N rate were included as fixed effects, and year was included as a random effect. The genotype × N rate interaction term was included when significant; otherwise, models containing only the main effects were used. Planned contrasts comparing each *pepck* mutant with W22 were performed separately within each N treatment using estimated marginal means (emmeans) with treatment-versus-control contrasts, with W22 specified as the control. Significance of model terms was assessed using type III analysis of variance with Satterthwaite’s method. Initial slopes of *A*_net_-*C*_i_ curves were compared using analysis of covariance (ANCOVA) of *A*_net_ at ambient CO_2_ concentrations from 0 to 200µmol mol^-1^ air. The maximum rate of CO_2_ assimilation was calculated as the maximum *A*_net_ value on the *A*_net_-Ci curve and compared among genotypes using a Student’s t-test. Mean *A*_net_ in the fluctuating light experiment was compared using two-way ANOVAs with genotype and light intensity as factors.

## Data Availability

All data supporting the findings of this study are provided in the main text and supplementary materials.

## Author Contributions

D.K.A., J.J.A., T.P.B., and S.P.M. conceived the research plans; A.S. performed the mutant screen; S.K., K.C., L.J., J.J.A. and D.K.A performed labeling and metabolite quantitation experiments; J.J.A. performed the field experiments based on S.P.M. methods; P.E. designed and performed physiological experiments based on A.B.C. methods. Supporting experimental analysis was performed by S.W., I.K., and K.M.M., who also performed supporting field studies and propagated the seed with M.L. M.L. developed and refined the PEPCK assays. J.Z. performed modeling and statistical analyses. J.J.A., D.K.A., T.L.M., and S.P.M. integrated and carefully evaluated data, with T.L.M. ’s contribution occurring prior to death on January 6, 2023. J.J.A. and D.K.A. drafted the manuscript with significant input from J.Z., K.M.M., and M.L., and additional contributions from all other authors. D.K.A. agrees to serve as the author responsible for contact and ensuring communication.

## Acknowledgements

This article is dedicated to the memory of Todd L. Mockler, a co-author with a passion for ideas that were ahead of his time, who passed away too soon and is greatly missed by the community. The authors thank the Plant Growth Facility (RRID:SCR_024902), Bioanalytical Chemistry Facility (RRID:SCR_001047), the Phenotyping Core Facility (RRID:SCR_019049), and the Subterranean Influences on Nitrogen and Carbon (SINC) Center at the Donald Danforth Plant Science Center. We thank the Roy J. Carver Biotechnology Center at the University of Illinois at Urbana-Champaign for sequencing RNAseq samples. Funding for the research was provided by the National Science Foundation Plant Genome Research Program postdoctoral fellowship NSF-1812235 to J.J.A. and NSF-2109387 to K.M.M., and through DBI-2242822, the United States Department of Agriculture National Institute of Food and Agriculture (2016-67013-24585, 2017-67013-26156, 2021-67013-33778, and 2023-67017-39419), National Institutes of Health (U01 CA235508). The United States Department of Agriculture Agricultural Research Service is acknowledged. This research was supported in part by the Office of Biological and Environmental Research in the DOE Office of Science (DE-SC0008769) to A.B.C. and (DE-SC0023142) to D.K.A. Support for the acquisition of the 6500 QTRAP LC–MS/MS was also provided by the National Science Foundation (DBI #1427621). The authors gratefully acknowledge additional technical help from Mrs. Sally R. Bailey and Drs. Saba Gill, Stewart Morley, Thiya Mukherjee, Mythri Jonnavittula, and Sarah Powers for help with fieldwork and biomass analysis. The authors also thank the three anonymous reviewers and the Editor for insightful questions and feedback that significantly improved the manuscript.

## Figure Legends

**Supplemental Table 1**: Significance values from mixed-effects model A.) for phenotypic traits and B.) for yield component traits.

**Supplemental Table 2:** List of qPCR primers.

**Supplemental Figure 1:** PEPCK activity measured by varying A.) the amount of PEP substrate or B.) the amount of biological supernatant containing the enzyme in both top and subtending leaves. Error bars represent mean ± standard error (n = 3 in A and n = 4 in B). Compact letter assignment represents significant differences (p < 0.05) among the three genotypes. No letter is displayed if there is no difference. (n = 4, ANOVA followed by Tukey’s multiple comparisons test).

**Supplemental Figure 2:** Total pool sizes in high- or low-N treatment for the molecules involved in the C_4_ shuttle: pyruvate (PYR), phosphoenolpyruvate (PEP), alanine (ALA), malate (MAL), and aspartate (ASP), or N metabolism: asparagine (ASN), glutamine (GLN), glutamate (GLU). Error bars represent mean ± standard error, n = 4.

**Supplemental Figure 3:** MIDs for ASP, MAL, PEP, PGA, and PYR at A) low N and B) high N conditions in W22 and *pepck1-2* lines. Error bars represent the mean ± standard error, n = 4.

**Supplemental Figure 4:** Average labeling for ASP, MAL, PEP, PGA, and PYR at A) low N and B) high N conditions in W22 and *pepck1-2* lines. Solid black curves represent GAM fit.

**Supplemental Figure 5:** Linear model fit to ASP and MAL on pool size scaled M+1 during 0-30s. Numbers on the plot are estimated slopes.

**Supplemental Figure 6:** Plant height in greenhouse-grown plants 55 days after sowing. A.) *pepck1-2* plants were visibly taller than W22 control plants at both high and low N B.) At high N, *pepck1-2* plants were 16.2 cm taller, at low N, plants were 14.1 cm taller (n = 40 plants; ***: p < 0.001, two-sided Welch’s test).

**Supplemental Figure 7:** Representative ears from pepck1*::Ds* and W22 control plants grown at high and low N in 2018. Supplemental Figure 7 provides images of field grown phenotypes and includes replicates in support of Figure 4.

**Supplemental Figure 8:** *A*_net_ was measured in fluctuating light environments in Pullman WA (A.) or St. Louis MO (B). Leaves were acclimated to high light (1500 μmol photons m^-2^ s^-1^), then they were exposed to alternating high and low light (250 μmol photons m^-2^ s^-1^) intensities in the following intervals: 1500, 250, and 1500 for ten minutes each then a series of fourteen one-minute intervals of low and high light intensities. Grey points represent W22 control plants; blue and green lines represent pepck1-1 and pepck1-2 mutant plants. Points represent means ± se for the light interval, n = 7-8 plants. Colored bands represent the mean during one-second measurement intervals. Supplemental Figure 8 includes replicates in support of Figure 5.

**Supplemental Data 1**: Biomass data for greenhouse grown plants.

**Supplemental Data 2**: Metabolite labeling from ^13^CO_2_ experiments.

**Supplemental Data 3**: Metabolite pool sizes for central intermediates.

**Supplemental Data 4**: Field data measurements for plants measured at R6 growth stage in 2016-2022.

