## Supplemental Figures for "Plant growth, biomass, and nitrogen utilization are impacted by mutations in C4 PEPCK1 in maize"

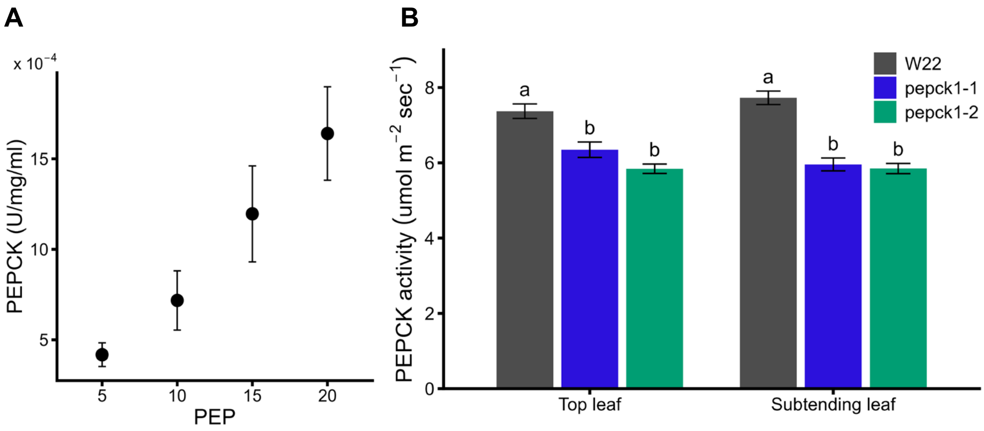


**Supplemental Figure 1:** PEPCK activity measured by varying A.) the amount of PEP substrate or B.) the amount of biological supernatant containing the enzyme in both top and subtending leaf. Error bars represent means ± standard error (n = 3 in A and n = 4 in B). Compact letter assignment represents significant differences ($p < 0.05$) among the three genotypes. No letter is displayed if there is no difference. (n = 4, ANOVA followed by Tukey's multiple comparisons test).

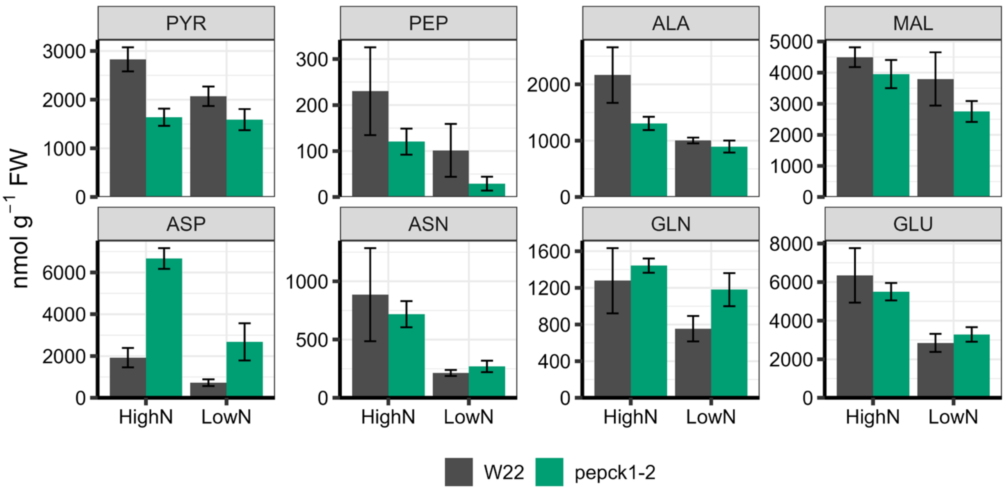


**Supplemental Figure 2:** Total pool sizes in high or low N treatment for the molecules involved in the C4 shuttle: pyruvate (PYR), phosphoenolpyruvate (PEP), alanine (ALA), malate (MAL), and aspartate (ASP) or N metabolism: asparagine (ASN), glutamine (GLN), and glutamate (GLU). Error bars represent means ± standard error, n = 4.

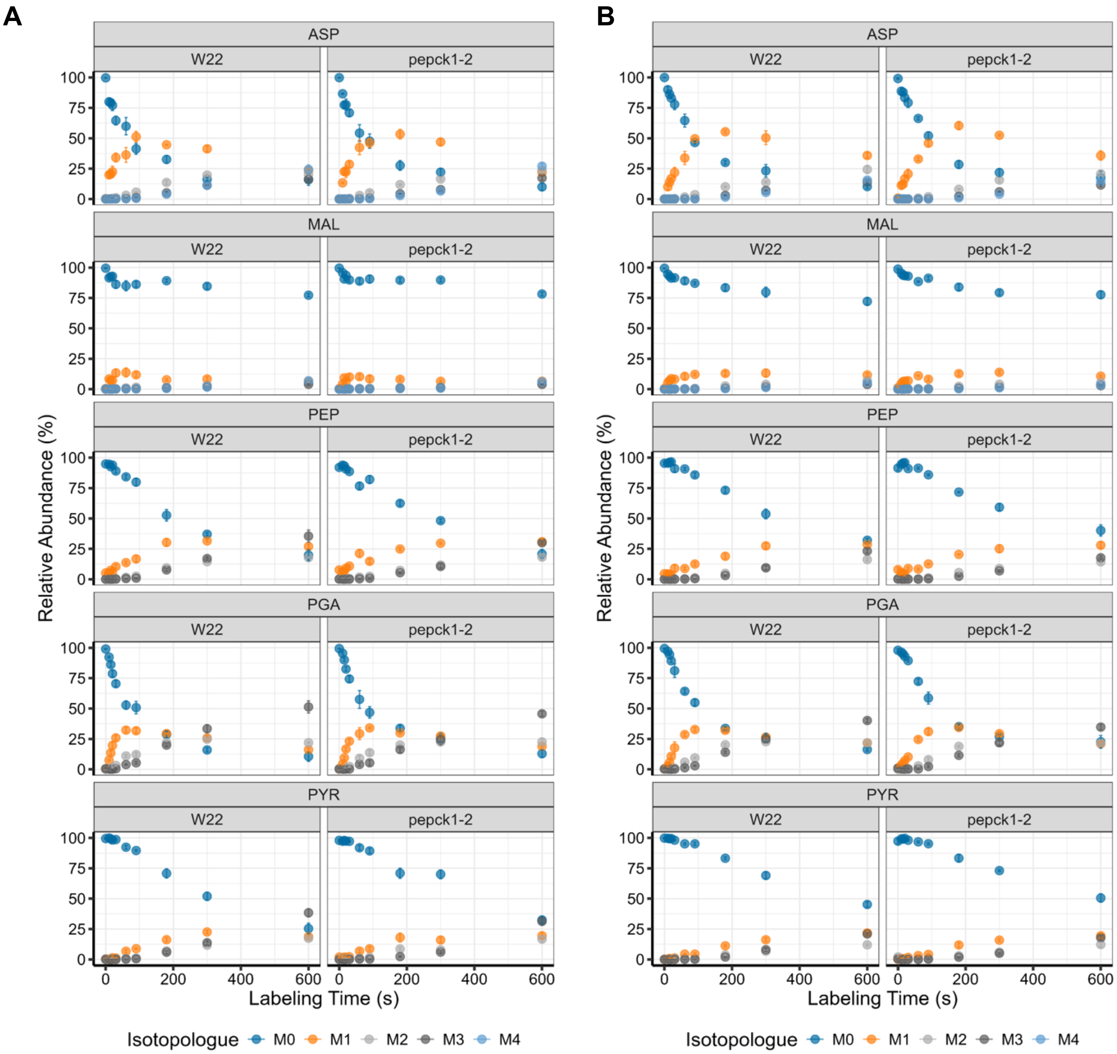


**Supplemental Figure 3:** MIDs for ASP, MAL, PEP, PGA, and PYR at A) low N and B) high N conditions in W22 and *pepck1-2* lines. Error bars represent means ± standard error, n = 4.

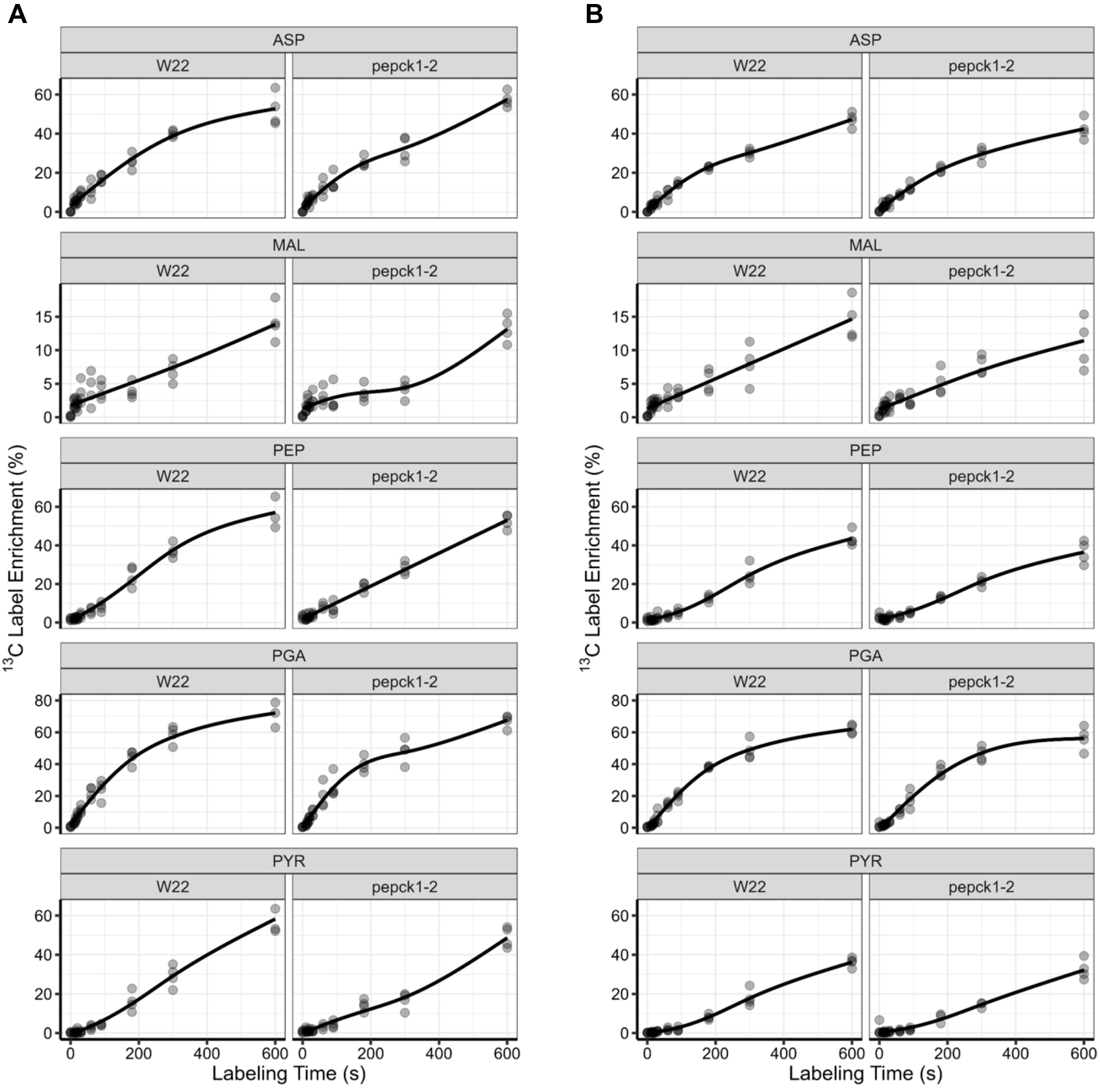


**Supplemental Figure 4:** Average labeling for ASP, MAL, PEP, PGA, and PYR at A) low N and B) high N conditions in W22 and *pepck1-2* lines. Solid black curves represent GAM fit.

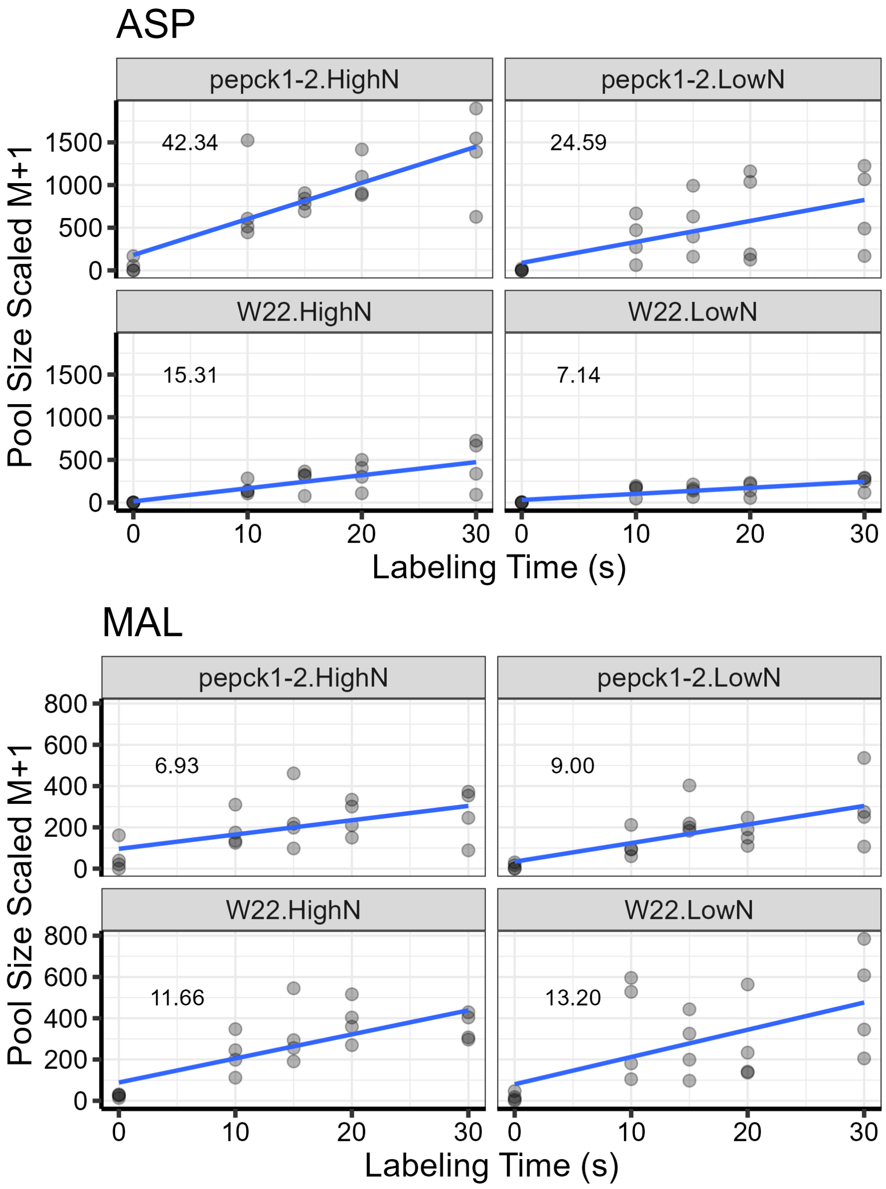


**Supplemental Figure 5:** Linear model fit to ASP and MAL on pool size scaled M+1 during 0-30s. Numbers on the plot are estimated slopes.

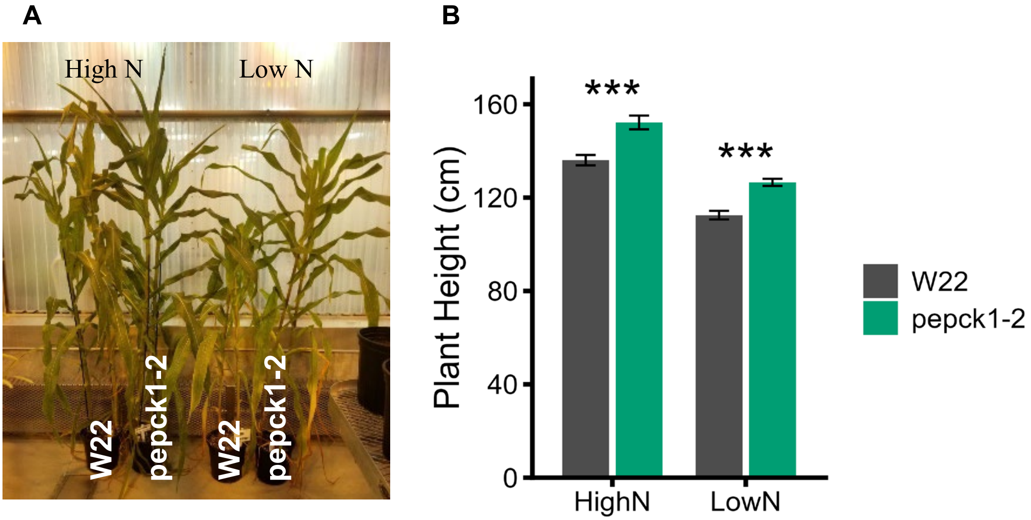


**Supplemental Figure 6:** Plant height in greenhouse-grown plants 55 days after sowing. A.) *pepck1-2* plants were visibly taller than W22 control plants at both high and low N B.) At high N, *pepck1-2* plants were 16.2 cm taller, at low N, plants were 14.1 cm taller. Error bars represent means ± standard error (n = 40 plants; ***: $p < 0.001$, two-sided Welch's test).

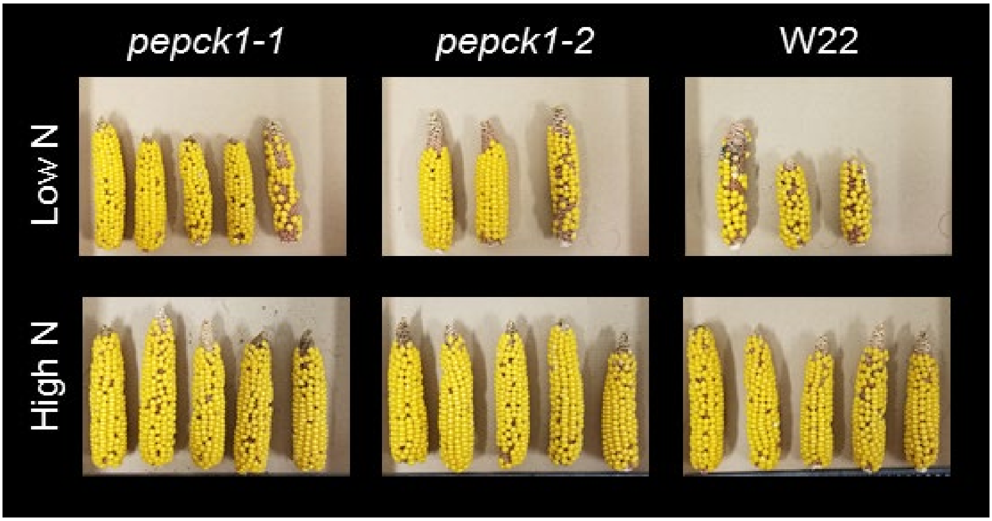


**Supplemental Figure 7**: Representative ears from pepck1*::Ds* and W22 control plants grown at high and low N in 2018. Supplemental Figure 7 provides images of field grown phenotypes and includes replicates in support of Figure 4.

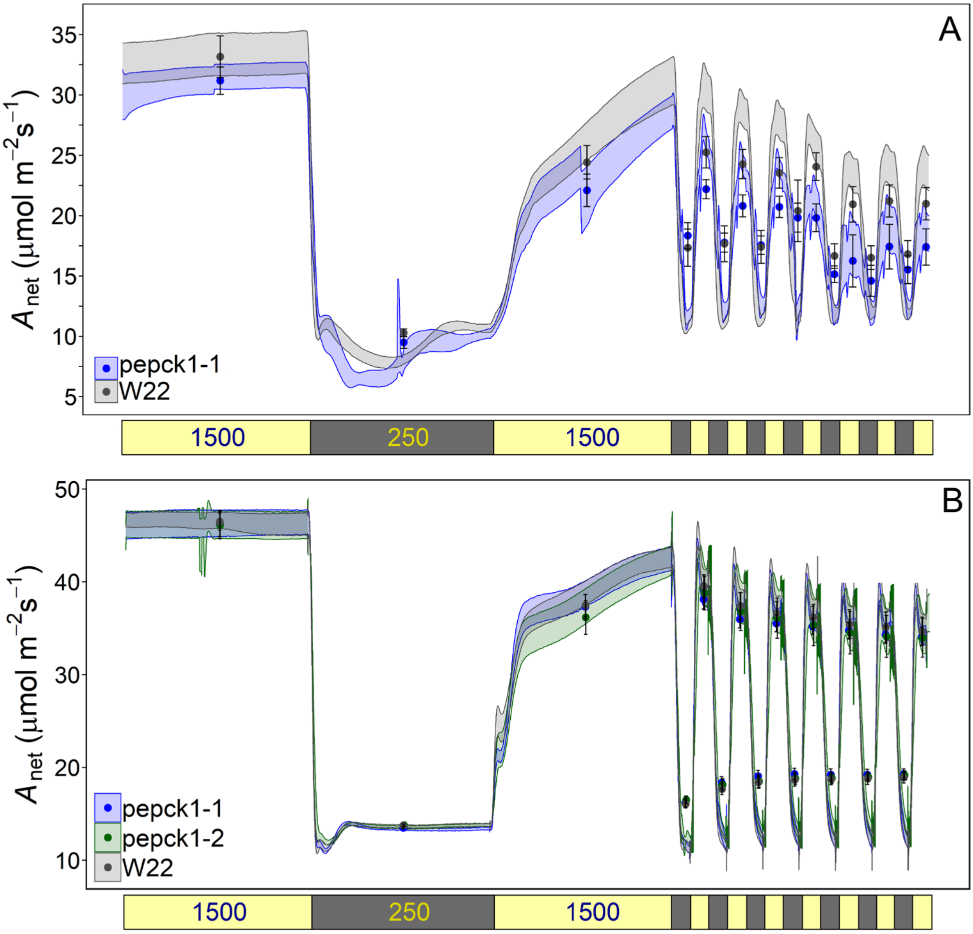


**Supplemental Figure 8:** $A_{net}$ was measured in fluctuating light environments in Pullman WA in the fall (A.) or St. Louis MO in the summer (B). Leaves were acclimated to high light (1500 µmol photons $m^{-2}$ $s^{-1}$), then they were exposed to alternating high and low light (250 µmol photons $m^{-2}$ $s^{-1}$) intensities in the following intervals: 1500, 250, and 1500 for ten minutes each then a series of fourteen one-minute intervals of low and high light intensities. Grey points represent W22 control plants; blue and green lines represent pepck1-1 and pepck1-2 mutant plants. Points represent means ± se for the light interval, n = 7-8 plants. Colored bands represent the means during one-second measurement intervals. Supplemental Figure 8 includes replicates in support of Figure 6.
