## Supplemental Tables for "Plant growth, biomass, and nitrogen utilization are impacted by mutations in C4 PEPCK1 in maize"

**Supplemental Table 1A: Significance values from mixed-effects model for phenotypic traits:** Significance for genotype and line were calculated using type III analysis of variance with Satterthwaite's method. Bold terms indicate significance at < 0.05.

| *Component* | *Plant Height* | *Cob Biomass* | *Stover Biomass* | *Stover N* | *Grain Biomass* | *Grain N* | *Total Biomass* | *Total N* | *Total NUtE* | *Grain NUtE* | *Harvest Index* |
| --- | --- | --- | --- | --- | --- | --- | --- | --- | --- | --- | --- |
| *Genotype* | **1.11e-08** | **0.034** | **0.044** | **0.040** | **8.44e-05** | **0.0004** | **0.0003** | **0.002** | 0.437 | **0.003** | **0.0003** |
| *N Rate* | **5.07e-06** | **3.09e-12** | **2.26e-13** | **<2e-16** | **1.17e-05** | **2.18e-07** | **7.8e-13** | **< 2.2e-16** | **< 2e-16** | **1.70e-6** | 0.803 |
| *Genotype x N Rate* | NA | **0.039** | NA | NA | NA | NA | NA | NA | NA | NA | NA |

**Supplemental Table 1B: Significance values from mixed-effects model for yield component traits:** Significance for genotype and line were calculated using type III analysis of variance with Satterthwaite's method. Bold terms indicate significance at < 0.05.

| *Component* | *Ear Length* | *Ear Width* | *Ear Weight* | *Kernel Row Number* | *Cob Weight* | *Seed Weight* | *100k weight* | *Seed Volume* | *100k volume* | *Kernel Number* | *Average Kernel Weight* | *Moisture* | *Protein* | *Starch* |
| --- | --- | --- | --- | --- | --- | --- | --- | --- | --- | --- | --- | --- | --- | --- |
| *Genotype* | **1.84E-06** | 0.954 | **0.009** | 0.511 | **8.99E-08** | **0.011** | 0.095 | **0.004** | 0.057 | **0.007** | 0.052 | 0.324 | 0.932 | 0.352 |
| *N Rate* | **< 2.2E-16** | **7.81E-09** | **2.98E-07** | **9.53E-07** | **< 2.2E-16** | **1.61E-04** | **9.61E-05** | **2.74E-06** | **4.64E-08** | **0.012** | **0.006** | **5.04E-08** | **4.81E-10** | **2.84E-05** |
| *Genotype x N Rate* | NA | NA | NA | NA | NA | NA | NA | NA | NA | NA | NA | **1.29E-4** | NA | NA |

**Supplemental Table 2: List of qPCR primers used.**

| qPCR Primer | Sequence (5’ -> 3’) |
| --- | --- |
| AS3_F | CCCTTACCTGCCCAAGCATA |
| AS3_R | CCTCCAGTTTATTGGCGCTGA |
| Asparaginase_F | ACAGAGAGAATCAAGCCGCC |
| Asparaginase_R | TCCGGTAGTCGAACAGGATG |
| AspAT-b_F | GCGCTGAAATCCAGAGGAAC |
| AspAT-b_R | AAAACTGCCCCACGAAGAGT |
| AspAT-m_F | TAAGCGTGGCATGGAAGTGT |
| AspAT-m_R | CGTCATGTTGTCACTCTGCG |
| CA1_F | CGCCCGACAAGTTCCACT |
| CA1_R | ACGCAGCTAATAAGTCCGGT |
| CA6_F | CTTGTAGAATGCCCGCCTTT |
| CA6_R | CGGTGTACTTGGTCTTGTCG |
| MDH3_F | AGGACCCAAACCCCAATTCC |
| MDH3_R | AACAGGGACATTCACATCGGT |
| PEPC_F | AAATCAGGAGGGCACAACCC |
| PEPC_R | GTGGCTCAAGGAACTCTTCG |
| PPDK_R | CCATGGAGCAGTTTGGAAGT |
| PPDK_F | TGGTGGCTGATGAGATAGCG |
| Housekeeping_NAC_F | CCGCCGTCAACAGGGAAATCTG |
| Housekeeping_NAC_R | GTAGCACGCCCAAGACCAACAG |
| Housekeeping_UPF_F | CACCCGGTTGGCTATGCTGTAC |
| Housekeeping_UPF_R | TGTGCTCCACCAGAAGGCTGAC |
